# Mechanisms of masking by Schroeder-phase harmonic tone complexes in the budgerigar (*Melopsittacus undulatus*)

**DOI:** 10.1101/2022.10.26.513930

**Authors:** Kenneth S. Henry, Yingxuan Wang, Kristina S. Abrams, Laurel H Carney

## Abstract

Schroeder-phase harmonic tone complexes can have a flat temporal envelope and either rising or falling instantaneous-frequency sweeps within periods of the fundamental frequency (F0), depending on the phase-scaling parameter C. Human thresholds for tone detection in a concurrent Schroeder masker are 10-15 dB lower for positive C values (rising frequency sweeps) compared to negative (falling sweeps), potentially due to the impulse response of cochlear filtering, though this hypothesis remains controversial. Birds provide an interesting animal model for studies of Schroeder masking because prior reports suggest less behavioral threshold difference between maskers with opposite C values. However, most behavioral studies focused on relatively low masker F0s, and neurophysiological mechanisms in birds have not been explored. We performed behavioral Schroeder-masking experiments in budgerigars (Melopsittacus undulatus) using a wide range of masker F0 and C values. The signal frequency was 2800 Hz. Neural recordings at the midbrain processing level characterized encoding of behavioral stimuli in awake animals. Behavioral thresholds increased with increasing masker F0 and showed minimal difference between opposite C values, consistent with prior studies. Neural recordings showed prominent temporal and rate-based encoding of Schroeder F0, and in many neurons, marked response asymmetry between Schroeder stimuli with opposite C values. Neural thresholds for Schroeder-masked tone detection were (1) in most cases based on a response decrement compared to the masker alone, consistent with prominent modulation tuning in midbrain neurons, and (2) generally similar between opposite masker C values. These results highlight the likely importance of envelope cues in behavioral studies of Schroeder masking.

## 1. Introduction

Schroeder’s algorithm can produce harmonic tone complexes (HTCs) with a flat temporal envelope and either upward-or downward-sloping instantaneous frequency sweeps within periods of the fundamental frequency (F0), depending on the polarity of the phase-scaling parameter C (Schroeder, 1970) (Fig. 1). Early on, it was discovered that HTCs generated with positive (downward-sweeping frequency) and negative (upward-sweeping frequency) C values differ in the extent to which they mask behavioral thresholds for pure-tone detection in humans, with lower thresholds by up to 20 dB for the positive C value compared to negative (Kohlrausch and Sander, 1995; Smith et al., 1986). Less masking by Schroeder HTCs with positive C values is not explainable by traditional power-spectrum models of masking (Fletcher, 1940) because Schroeder HTCs generated with opposite C values have identical magnitude spectra; these signals share the same waveform when one signal is time reversed.

**Fig. 1.**
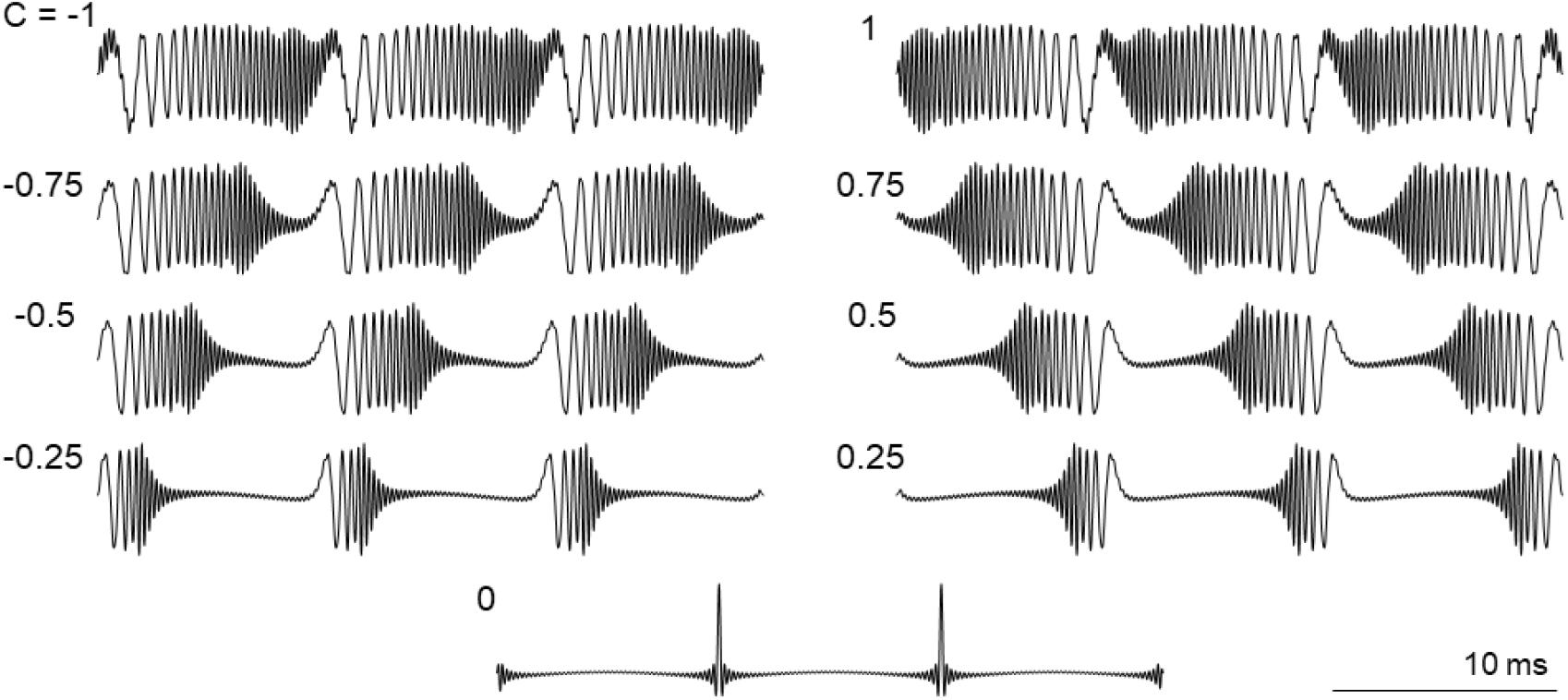
Waveforms of Schroeder-phase harmonic tone complexes (HTCs) produced with the same fundamental frequency (F0; 100 Hz) and values of the phase-scaling parameter, C, ranging from -1 to 1. Positive C values (right column) produce downward-sloping instantaneous frequency sweeps within F0 periods whereas negative C values (left column) produce upward-sloping frequency sweeps. Lower absolute C value decreases the duty cycle of the waveform, with C of zero (bottom panel; all components in cosine phase) producing an impulse train.

The underlying reason for masking differences between Schroeder C polarities remains uncertain. One hypothesis is that the difference originates from asymmetry of mechanical filtering by the mammalian cochlea (Smith et al., 1986). Specifically, recordings of basilar membrane motion in response to clicks show that the impulse response of the cochlea exhibits an upward-sweeping instantaneous frequency glide in response to clicks – a property associated with asymmetry in the shape of pure-tone frequency tuning curves (Recio et al., 1998; Strube, 1985). Filters with an upward-gliding impulse response are expected to produce more prominent response envelope fluctuations (i.e., “peakier” temporal-response profiles) for Schroeder HTCs with positive C values compared to negative. The deeper and longer low-amplitude periods in the cochlear response to maskers with positive C value could potentially provide opportunities for detection of an added low-level tone, thereby leading to the observed lower masked threshold for the positive-C condition.

Animal models have been used to test the hypothesized contribution of cochlear filtering asymmetries to Schroeder-masking differences between opposite C polarities, but so far have yielded an incomplete understanding of underlying mechanisms. Studies in chinchilla and guinea pig recorded basilar membrane motion in response to Schroeder HTCs to test for the predicted difference in response envelope fluctuations between opposite C polarities (Recio and Rhode, 2000; Summers et al., 2003). Results were partly consistent with an explanation based on cochlear impulse responses, showing the expected larger fluctuations for responses to Schroeder HTCs with positive C values compared to negative. However, measurements were restricted to basal cochlear regions with characteristic frequencies (CFs) above the frequency range of target signals typical of human behavioral studies. Note that for CFs below 1 kHz, the mammalian cochlea appears to transition to downward-sweeping frequency glides of the impulse response based on auditory-nerve-fiber recordings (Carney et al., 1999; Recio-Spinoso et al., 2005). Filters with downward-gliding impulse responses should theoretically produce a peakier response to Schroeder HTCs with negative C values rather than positive, and consequently less masking for the negative-C condition. However, less masking by Schroeder HTCs with negative C values has not been observed in any human behavioral study (Kohlrausch and Sander, 1995; Smith et al., 1986). While it remains possible that chinchillas or guinea pigs show lower thresholds for the negative-C Schroeder masking condition at test frequencies below 1 kHz, animal behavioral studies of Schroeder masking have not been conducted in any other nonhuman mammal to our knowledge. Consequently, it remains unclear whether asymmetry of cochlear filtering can explain observed behavioral masking differences between Schroeder C polarities.

Birds provide an interesting animal model for studies of Schroeder masking because many species use vocalizations containing rapid frequency changes as part of their social communication system and can perform complex auditory detection and discrimination tasks (Dooling et al., 2000). In contrast with human results, the few behavioral studies of Schroeder masking in birds have found minimal threshold differences for tones masked by Schroeder HTCs with opposite C values. In budgerigars, thresholds were determined for detection of 1, 2.8, and 4-kHz tones masked by Schroeder HTCs with positive or negative *C* values, for comparison with human thresholds determined using the same stimuli (Leek et al., 2000). Whereas human subjects showed the expected lower threshold of up to 15-20 dB for positive-C Schroeder maskers compared to negative C values, budgerigar thresholds were either similar between opposite C values or slightly lower for maskers with negative C values. In two follow-up studies, budgerigars, zebra finches, and canaries were tested across a wider range of stimulus conditions including higher F0s up to 400 Hz (Dooling et al., 2001), and multiple, intermediate C values between -1 and +1 (Lauer et al., 2006). Note that lower absolute C values produce stimuli with reduced duty cycles and, consequently, faster instantaneous frequency sweeps within F0 periods, with C of zero producing an impulse train associated with all components in cosine phase (Fig. 1). Even over this wider range of conditions, which included faster rates of instantaneous frequency change that were potentially better suited to the mechanics of the avian inner ear, birds showed little appreciable difference in masking between Schroeder HTCs generated with opposite C values (Dooling et al., 2001; Lauer et al., 2006).

The reason for less masking difference between Schroeder HTCs of opposite C polarity in birds compared to humans is unclear due to limited physiological investigations. One possibility is that the avian cochlea differs from that of mammals in its impulse response, perhaps lacking a prominent instantaneous frequency glide [see barn owl auditory-nerve recordings potentially consistent with this interpretation (Fontaine et al., 2015)], or perhaps showing a glide that is faster than the rate of frequency change found in Schroeder maskers from existing behavioral studies. Despite basic physiological similarities between birds and mammals in auditory-nerve frequency tuning, phase-locking limits, and other response properties of auditory-nerve fibers (Manley et al., 1985; Sachs et al., 1974), the sensory epithelium in birds is considerably shorter and wider than in mammals, with dozens of hair cells spanning its width in apical low-CF regions (Takasaka and Smith, 1971). If these anatomical differences result in a cochlear impulse response without a prominent frequency glide (see Fontaine et al., 2015), this could explain less masking difference between Schroeder HTCs of opposite polarity. While Dooling and colleagues found stronger compound auditory-nerve action potentials for negative-C Schroeder HTCs in several bird species (Dooling et al., 2002), these gross potentials are strongly influenced by cross-channel neural synchrony and are therefore not directly relatable to current hypotheses for human masking asymmetry, based on a single-channel framework.

To gain further insight into mechanisms of Schroeder masking, and into possible reasons for minimal Schroeder masking asymmetry in birds, we performed parallel behavioral and neurophysiological experiments in budgerigars using tones masked by Schroeder HTCs. In the first experiment, behavioral studies were conducted in animals trained using operant-conditioning procedures over a wide range of F0 and C values, to improve the possibility of detecting masking differences between Schroeder C polarities if they exist. To expand upon prior studies, masking conditions included previously untested C values of ±0.5 at F0s of 200 and 400 Hz. Furthermore, stimuli were presented using a roving-level paradigm for which the overall masker level was randomly selected on each trial from a 20-dB range centered on 80 dB SPL (a typical level used in prior studies), thereby diminishing the extent to which overall loudness and/or single-channel energy cues could be used to perform the task. In the second neurophysiological experiment, we made recordings in awake passively listening animals of extracellular activity in the central nucleus of the avian inferior colliculus (IC), which is a large and nearly obligatory midbrain processing center in the ascending auditory pathway. The IC shows pronounced temporal and rate-based encoding of acoustic-envelope fluctuations at amplitude-modulation frequencies up to several hundred Hz (Henry et al., 2017a; Joris et al., 2004), encompassing the range of envelope frequencies found in Schroeder HTCs. Moreover, a recent study of gerbil IC revealed neural selectivity for C polarity of Schroeder HTCs based on average discharge rate and spike timing (Steenken et al., 2022). The present study obtained budgerigar IC neural responses to Schroeder HTCs across widely ranging F0 and C values and evaluated the changes in neural activity that occur upon addition of a tone increment to Schroeder maskers, providing insight into potential neural cues used by animals to perform the behavioral masked-detection task.

## 2. Methods

### 2.1. Behavioral experiments

Behavioral experiments were conducted in four adult budgerigars of either sex (2 male; 2 female) using previously described operant conditioning procedures (Henry and Abrams, 2021; Henry et al., 2017a; Henry et al., 2020; Henry et al., 2017b; Henry et al., 2016; Wang et al., 2021; Wong et al., 2019). All procedures were approved by the University of Rochester Committee on Animal Resources.

#### 2.1.1. Stimuli

Stimuli in the behavioral experiments were Schroeder-phase HTCs presented with or without a simultaneously-gated 2.8-kHz tone signal. Schroeder HTCs were generated with component phases scaled according to Schroeder’s (1970) algorithm,

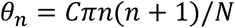

where *θ*_*n*_ is the phase of the *n*th harmonic, *N* is the total number of harmonics, and *C* is a scaling parameter. Harmonic components ranged in frequency from 2F0 to 5000 Hz and were presented with equal level. The overall level of the masker was 80 dB SPL prior to application of random level variation for some conditions (i.e., roving-level paradigm; see below). Consequently, for F0s of 100, 200, and 400 Hz, the number of components included in the Schroeder masker was 49, 24, and 11, respectively, and the level of individual masker components prior to roving was 63.10, 66.20, and 69.59 dB SPL, respectively.

Schroeder maskers and pure-tone targets were simultaneously gated on target trials (50% of all trials) and had a duration of 260 ms, including 20-ms raised-cosine onset and offset ramps. The phase of the 2.8-kHz pure tone target was equal to that of the masker frequency component at the same frequency. The masker alone was presented on standard trials. Threshold signal-to-masker ratios for tone detection are expressed in dB relative to the level of single masker components, as in previous studies (Kohlrausch and Sander, 1995; Lauer et al., 2006; Smith et al., 1986).

The first behavioral experiment assessed Schroeder masking in three animals over a wide range of masking conditions using F0s of 100, 200 and 400 Hz and C values of 0, ±0.5, and ±1. Testing was conducted using a roving-level paradigm for which the overall stimulus level on each trial was randomly varied with respect to the mean level based on a uniform distribution spanning -10 to +10 dB with 1-dB resolution. A second behavioral experiment compared Schroeder-masked thresholds between fixed-level and roving-level test conditions in four animals, using F0 of 100 Hz and C values of 0 and ±1. Testing alternated between fixed- and roving-level conditions within tracking sessions, as described in detail below, to assess the effect of the roving-level paradigm.

#### 2.1.2. Apparatus

Behavioral experiments were conducted in four identically constructed small-animal acoustic isolation chambers (0.3 m^3^) lined with sound-absorbing foam. Chambers contained a house light, overhead loudspeaker (Polk Audio MC60; Baltimore, MD), seed dispenser (ENV-203 mini; Med Associates, St. Albans, VT), feeding trough, small wire-mesh cage with a perch, and three illuminated response switches. The loudspeaker was positioned 20 cm above the location of the animal’s head during observing responses (see below). Behavioral tests were controlled using custom MATLAB software (The MathWorks, Natick, MA) running on a PC. Stimulus waveforms were generated digitally with 50-kHz sampling frequency and converted to analog signals (National Instruments PCI-6151; Austin, TX) before power amplification (Crown D-75A).

Custom-built electronic equipment was used to process input from the response switches and control the house light and seed dispenser. Stimuli were calibrated using a pre-emphasis filter generated based on the output of a ½” precision microphone (Brüel and Kjær type 4134, Marlborough, MA) in response to 249 log-spaced tone frequencies from 50-15,100 Hz. The pre-emphasis filter corrected the stimulus spectrum for the frequency response of the loudspeaker system prior to analog conversion.

#### 2.1.3. Procedure

Budgerigars were trained using operant-conditioning procedures to perform a single-interval, two-alternative, auditory discrimination task. Birds initiated a trial by pecking the center observing-response switch, resulting in presentation of a single stimulus. The stimulus was either a standard Schroeder HTC masker or target with a tone signal added to the masker. Standard and target stimuli were presented with equal probability of 0.5. The correct response to the standard stimulus was the right switch and the correct response to the target stimulus was the left switch. Animals had 3 seconds to report a response following stimulus onset. Correct responses were reinforced with delivery of hulled millet seeds, whereas incorrect responses resulted in a 5-s timeout during which the house light was turned off. The trial was aborted in rare instances for which animals did not report a response (<0.2% of trials). The number of millet seeds was adaptively varied (1 or 2 seeds) using an automated procedure to control response bias. Bias was calculated based on the most recent 50 trials as 0.5 times the sum of the Z scores of the hit and false-alarm rates (Macmillan and Creelman, 1991). Bias was controlled to stay near zero by delivering additional seeds for correct responses made with the switch the animal was biased against. Testing was conducted 6-7 days per week during morning and afternoon blocks separated by four hours, and consisting of several hundred trials each.

Animals were initially trained to behaviorally discriminate the target stimulus from the standard at a relatively high signal level of 10 dB re: masker component level. Once 90% correct discrimination performance was attained, masked thresholds were tested repeatedly using two-down one-up adaptive-tracking procedures (Levitt, 1971). Behavioral-tracking sessions began at a signal level of 10 dB re: masker component level. The signal level was reduced following every two consecutive correct responses to target trials (hits) with the same signal level, and increased following every incorrect response to a target trial (miss). Responses on standard trials (false alarms and correct rejections) were reinforced but did not affect the signal level of the track. The step size of the tracking procedure was varied based on the number of reversals (upward to downward or vice versa) of the signal-level track across target trials. The initial step size of 3 dB was decreased to 2 dB after 2 reversals and to 1 dB after 4 reversals. Tracks continued for at least 15 reversals until the signal level of the final eight track reversal met two stability criteria. Both (1) the standard deviation of the final eight reversal values and (2) the difference between the means of the first and second sets of four reversals in the final eight were required to be less than 3 dB (i.e., the track had to be consistent, without an upward or downward trend of the final reversal points). Tracks for which the overall absolute bias exceeded 0.3 were excluded from further analysis.

For the first experiment, the mean of the final eight reversal values was taken as the track threshold. For the second experiment, extended-duration tracks (Wang et al., 2021) were used to evaluate the effect of roving stimulus level on Schroeder-masked tone detection within sessions, and fewer tracks were completed daily. For extended tracks, the masker level remained fixed at 80 dB SPL until the stability criteria described above were satisfied. This initial portion of the track provided reversal points for calculation of the fixed-level masked threshold. Thereafter, the track was continued (without resetting the signal level) for an additional 10 reversals under the roving-level condition (±10 dB across-trial variation in masker level with respect to the fixed masker level from the first part of the track) until the track stability criteria were once again satisfied. This second portion of the track was used to calculate the roving-level threshold (based on the final eight reversal points as above).

For both experiments, tracking sessions were conducted repeatedly on the same stimulus condition until at least 13 unbiased thresholds (or threshold pairs for experiment 2) were obtained, with absolute response bias less than 0.3, and two across-track stability criteria were satisfied. Both (1) the standard deviation of the final six unbiased track thresholds and (2) the mean difference between the first and second half of the final six unbiased thresholds were required to be less than 3 dB. For both experiments, stimulus conditions were tested in different random sequences across animals. Animals were tested 2-4 times on each condition, with each testing period separated by testing on the other conditions.

#### 2.1.4. Analysis

The last 25 unbiased-track thresholds were averaged to calculate masked thresholds of individual animals on each condition. Thresholds were analyzed in R (v. 4.1.3) using linear mixed-effect models (Bates et al., 2015). Models incorporated random intercepts for each animal subject, to account for repeated observations, and included fixed effects of F0, C, and roving level (for experiment two). Two-way interactions were included and dropped when not significant (p > 0.05) in order of decreasing p value. Degrees of freedom for post-hoc t-tests were calculated based on the Satterthwaite approximation.

### 2.2 Neurophysiological recordings

Neurophysiological recordings were made from the IC in four adult budgerigars of either sex (two male; two female) using previously reported procedures (Henry et al., 2017a; Henry et al., 2016; Wang et al., 2021). Different animals were used for neural recordings than those from the behavioral experiments. All procedures were approved by the University of Rochester Committee on Animal Resources.

#### 2.2.1 Neural implantation procedure

A probe assembly consisting of 1-2 microelectrodes mounted to a miniature microdrive was implanted into the right IC under anesthesia using a stereotaxic approach. Microelectrodes were either single pure iridium wires or four-channel silicon probes with impedance ranging from 1.5-3 MΩ. Anesthesia was induced with subcutaneous injection of ketamine (3-5 mg/kg) and dexmedetomidine (0.08-0.1 mg/kg), and maintained throughout the 2-3 h implantation surgery with slow subcutaneous infusion of the same agents (ketamine: 6-10 mg/kg/h; dexmedetomidine: 0.16-0.27 mg/kg/h) in lactated ringers solution. When possible, electrodes were initially implanted superficially in the dorsal region of the IC tuned to low frequencies. This method allowed sampling of response regions tuned to higher CFs by advancing the electrodes deeper into the IC across recording sessions with the microdrive control screw. Electrodes were typically advanced 30 um between recording sessions. In total, recordings were made from 142 multi-unit sites and 8 sites with well-isolated single-unit activity (criteria described below).

#### 2.2.2. Apparatus

Neurophysiological recordings were made with animals perched in a small wire-mesh cage located inside a walk-in acoustic isolation chamber. The chamber was lined with sound-absorbing foam and contained a loudspeaker (MC60, Polk Audio) positioned 45 cm from the normal location of the animal’s head in the same horizontal plane. A closed-circuit video system was monitored to ensure that animals remained perched and facing the loudspeaker throughout recording sessions.

Stimuli were generated in MATLAB with a sampling frequency of 50 kHz and converted to analog at full scale using a data acquisition card (±10V; PCIe-6251, National Instruments). Signals were then attenuated to the specified level (PA5; Tucker-Davis Technologies) followed by power amplification (D-75A, Crown Audio). Stimulus calibration was accomplished with a digital “pre-emphasis” filter that compensated for the frequency response of the system. The filter was designed based on the output of a 0.25” precision microphone (type 4938, Brüel & Kjær) placed at the location of the animal’s head in response to 249 log-space tone pips ranging in frequency from 0.05-15.1 kHz.

Neural recordings were made using a multi-channel recording system (RHD2132 amplifier chip and C3100 USB interface board, Intan Technologies), referenced to one of the anchor screws for the microdrive assembly. Recordings were high-pass filtered at 150 Hz and sampled on the amplifier chip at 30 kHz for storage on the computer hard drive. In post processing, recordings were resampled at 50 kHz and band-pass filtered from 0.75-10 kHz (1000-point finite impulse response) to reduce local field potentials. Thereafter, a third-order Teager-energy operation (Choi et al., 2006; Wang et al., 2021) was applied for spike detection based on a visually determined threshold criterion. Sessions with consistent spike shape and spike amplitude through the recording period and less than 1% of inter-spike intervals less than 1 ms were classified as single-unit sessions. Both single- and multi-unit recordings were analyzed, as described below.

#### 2.3.3. Stimuli

Stimuli used in neurophysiological recordings were pure tones for determination of the frequency response map, sinusoidally amplitude-modulated (SAM) tones, Schroeder-phase HTCs, and Schroeder-phase HTCs with an added tone. The frequency response map shows average discharge rate as a function of stimulus frequency and level. Tone stimuli for the response map ranged in frequency from 0.25-8.0 kHz with 8 steps per octave, and were presented at levels from 15-75 dB SPL in 10-dB steps. Tone duration was 100 ms including 10-ms raised-cosine onset and offset ramps. Stimuli were presented in random sequence with three repetitions of each frequency-level combination, and 250-ms of silence between successive stimuli.

SAM tones were used to characterize modulation tuning of each recording site with a modulation-transfer-function (MTF) approach. The MTF shows average discharge rate as a function of stimulus modulation frequency. Stimuli had modulation frequencies ranging from 4 to 1024 Hz with 3 steps per octave and carrier frequency equal to the estimated CF. Overall level was 65 dB SPL and the depth of modulation was 0 dB (100%), except for the case of an unmodulated CF tone also included in the stimulus set. Stimulus duration was 800 ms including 50-ms raised-cosine onset and offset ramps. MTF stimuli were presented in random sequence for 5 or 10 repetitions with 350-ms of silence between stimuli.

Responses to Schroeder HTCs were obtained for F0s of 50, 100, 200, and 400 Hz and C values from -1 to +1 with a step size of 0.25. Harmonic frequency components between 250 Hz and 5000 Hz (inclusive) were included in the stimulus, and overall level was scaled to 60 dB SPL. Stimuli were 400 ms in duration, including 25-ms raised-cosine onset and offset ramps, presented for 20 repetitions in random sequence with 200 ms of silence between stimuli.

Responses to Schroeder-maskers with an added tone signal were obtained for masker F0s of 100, 200, and 400 Hz and C values from -1 to +1 with a step size of 0.5. Frequency components between 200 Hz and 5000 Hz (inclusive) were included in the masker, tone frequency was equal to that of the nearest masker component to CF, and tone phase was equal to that of the masker component at the same frequency. Tone and masker were simultaneously gated with a duration of 260 ms, 20-ms onset and offset ramps, and 250 ms of silence between stimuli. Signal level re: single masker components ranged from -33 to 0 dB, spanning the observed range of behavioral thresholds, and included a masker with no added tone signal to allow estimation of masked neural detection thresholds (see below). The overall level of the masker was 60 dB SPL. Stimuli were presented in random sequences, with 20 repetitions of each stimulus condition.

#### 2.3.4. Analyses

CF was determined from frequency response maps as the stimulus frequency at which the neural response was most sensitive (i.e., responded at the lowest sound level). Best modulation frequency was defined as the modulation frequency of the MTF peak, and the amplitude of modulation tuning was estimated as the percent difference between the peak MTF rate and the response rate for the unmodulated stimulus.

Neural responses to Schroeder HTCs were quantified with both an average-rate analysis and a temporal metric calculated as the peak-to-peak amplitude of single-trial period histograms. The temporal metric quantifies the mean amplitude of instantaneous rate fluctuations at F0 and is hereafter called “rate fluctuation” for brevity. Rate fluctuation was calculated after first integrating each single-trial spike-train response over a duration equal to 1/25^th^ of the F0 period for F0s up to 200 Hz (50 Hz: 0.8 ms; 100 Hz: 0.4 ms; 200 Hz: 0.2 ms) and over 0.2 ms for F0 of 400 Hz. Schroeder bias was calculated for both average rate and rate fluctuation as the mean response difference between positive and negative Schroeder polarities divided by the pooled standard deviation. Hence, positive (negative) values indicate a greater response amplitude for the positive (negative) Schroeder stimulus and the magnitude of bias metric relates to separation between the means in terms of standard deviation, as in a d-prime analysis (Egan, 1975).

Neural thresholds for detection of a CF tone added to a Schroeder HTC were estimated from neurophysiological data using receiver-operating characteristic (ROC) analysis (Egan, 1975). For each tone level presented, the discrimination performance of the neuron was calculated as the area under the ROC curve between the response distribution observed for the masker alone and the distribution for the tone-plus-masker stimulus. Area under the ROC curve corresponds to the percent separation (100 – percent overlap) between the response distributions. Response distributions were based on 20 stimulus repetitions. The function describing discrimination performance across signal levels was linearly interpolated with 0.1-dB resolution and inverted (i.e., 100 – discrimination performance) in cases for which performance at the highest signal-to-masker ratio was less than 50% correct. Threshold was defined as the highest signal-to-masker ratio above which discrimination performance consistently exceeded 70.7% correct, which is the average correct performance of an unbiased theoretical observer performing a two-down one-up track (Levitt, 1971).

## 3. Results

### 3.1. Behavioral Results

#### 3.1.1. Effects of F0 and C on Schroeder-masked thresholds

Budgerigars were trained with operant-condition procedures to detect a 2.8-kHz tone added to a Schroeder HTC masker. Masked detection thresholds were tested in three animals using two-down one-up tracking procedures (Fig. 2A) and a ±10-dB roving-level paradigm that reduces the utility of single-channel energy cues for performing the task. Maskers had F0 of 100, 200, or 400 Hz and C values of -1, -0.5, 0, +0.5, or 1 (15 conditions). Each animal completed the experiment over a period of 8-9 months consisting of 850-950 test sessions.

**Fig. 2.**
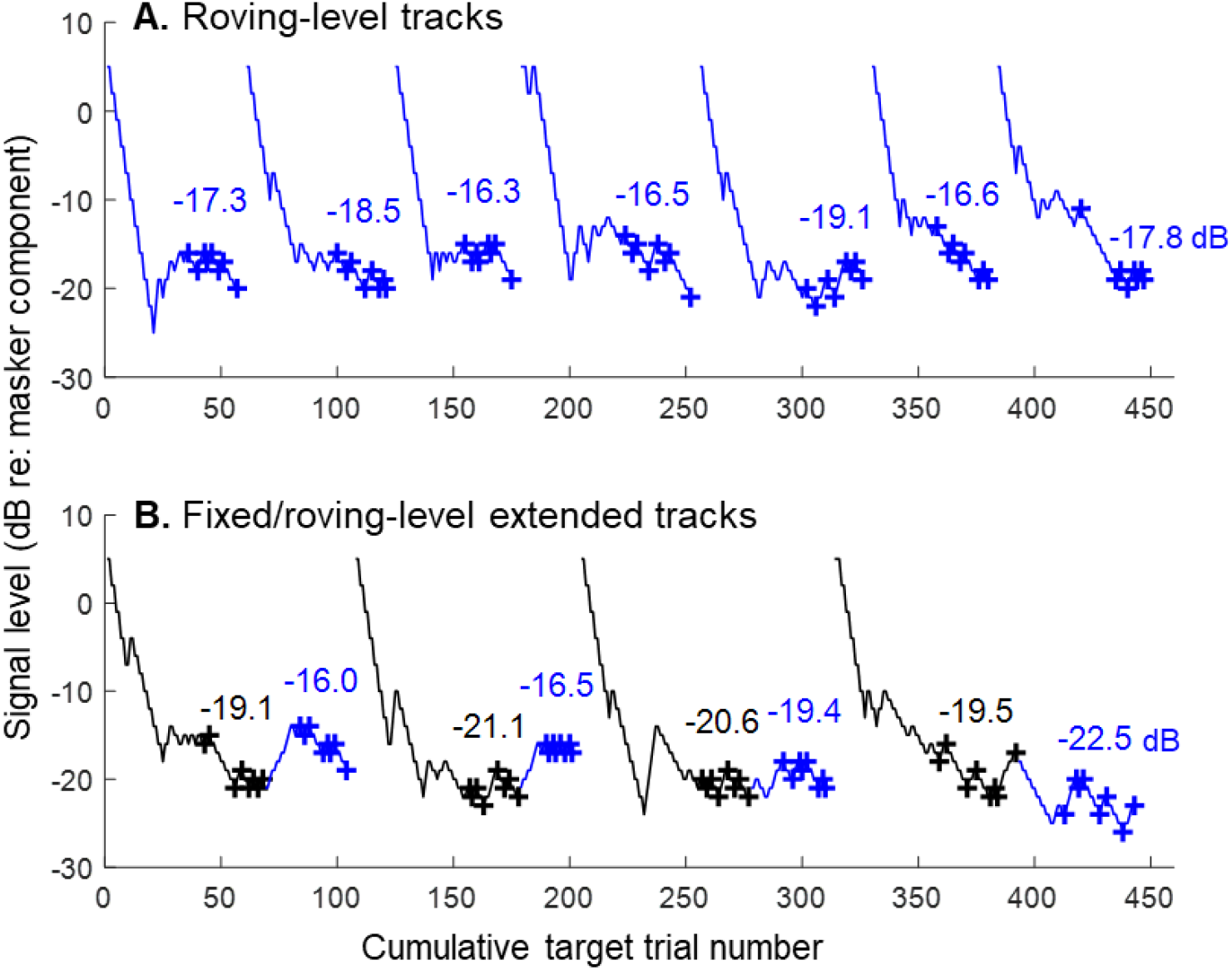
Representative results of two-down one-up tracking sessions in trained budgerigars. Tracks show variation in signal level across trials.. Roving-level tracks (A) are obtained with ±10-dB random variation in stimulus level across trials that reduces the utility of single-channel energy cues for performing the task. Extended tracks (B) transition from a fixed-(black) to the roving-level (blue) test procedure within sessions. Crosses indicate final reversal points used to calculate track thresholds, given above each track in dB. F0 is 100 Hz and C is +1.

Schroeder-masked behavioral thresholds of individual animals were obtained for each condition (Fig. 3A) based on the final 25 unbiased tracks. Masked thresholds increased for higher F0 values and for greater absolute C values, with C values of ±0.5 producing intermediate thresholds between C of zero, for which thresholds were lowest, and C of ±1, for which thresholds were highest. A mixed-effects model analysis of masked thresholds showed significant effects of F0 (F_2,28_=34.14, p<0.0001), C (F_4,28_=17.59, p<0.0001), and the F0-by-C interaction (F_8,28_=2.76, p=0.0220). The interaction was due to greater elevation of thresholds with increasing F0 for C of zero and ±0.5 compared to ±1. No significant differences in threshold were observed between negative and positive polarities of the same C value at F0s of 100 Hz (C ±1: -1.77 ±1.85 dB; t_28_=-0.96, p=0.35; C ±0.5: -2.55 ±1.85 dB; t_28_=-1.38, p=0.18), 200 Hz (C ±1: 2.02 ±1.85 dB; t_28_=1.09, p=0.28; C ±0.5: 1.39 ±1.85 dB; t_28_=0.75, p=0.46), or 400 Hz (C ±1: 1.12 ±1.85 dB; t_28_=0.60, p=0.55; C ±0.5: -0.46 ±1.85 dB; t_28_=-0.25, p=0.80; least-squared mean difference ±1 SE and pairwise comparisons of least-squared means). These results indicate no asymmetry of tone masking by Schroeder HTCs in the budgerigar.

**Fig. 3.**
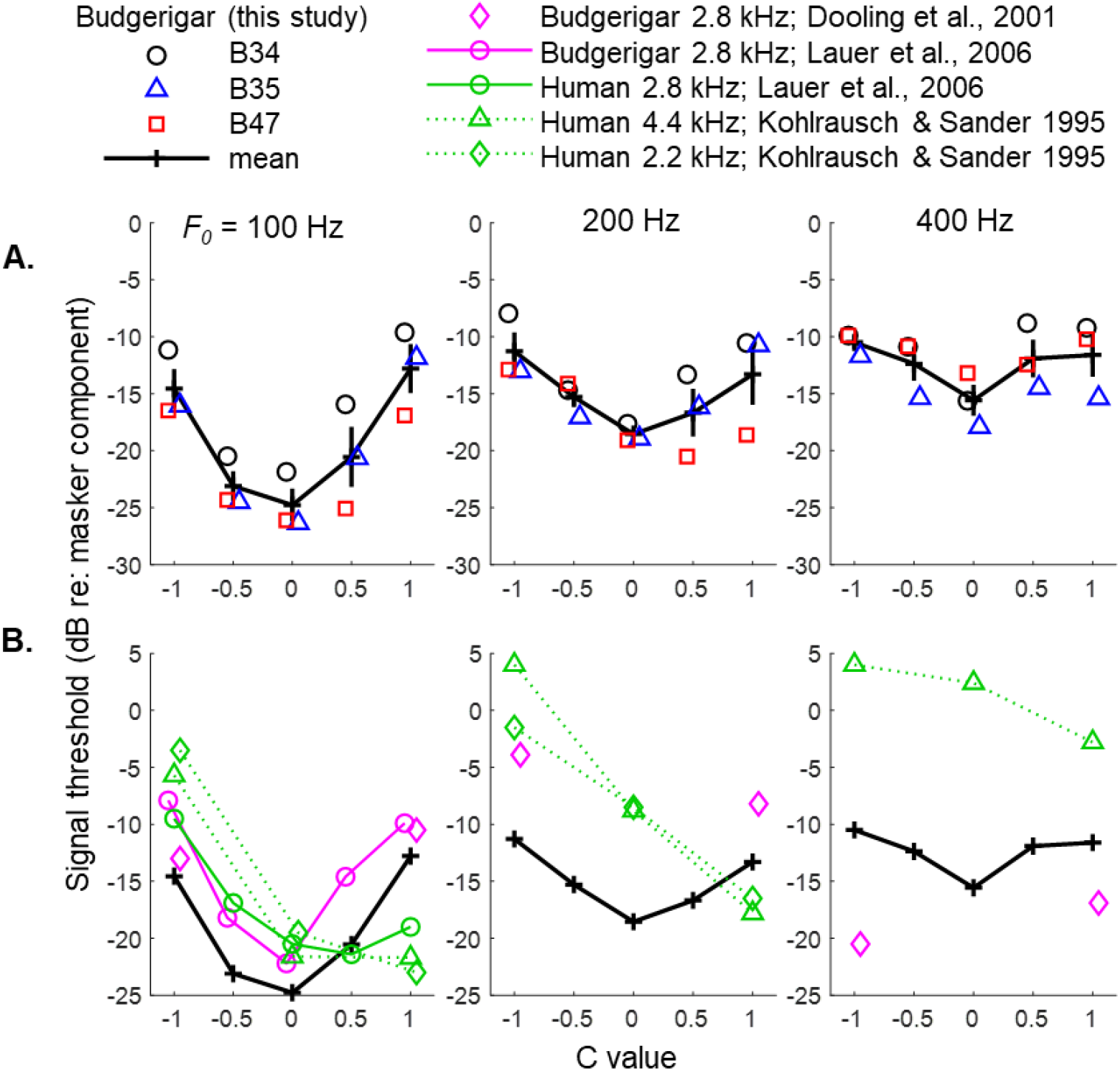
Behavioral thresholds of individual budgerigars for detection of a tone signal masked by a Schroeder HTC (A). Trend lines and error bars are means and across-subject standard deviations, respectively. Budgerigar Schroeder-masked tone thresholds increase for higher F0s, increase for higher absolute C values, and are similar between opposite polarities of the same C value. Mean thresholds from the current study compared to previously published results from humans and other budgerigar studies (B). Budgerigars show less masking difference between positive and negative C value than humans.

Several previous studies quantified Schroeder masking in budgerigars and humans using similar stimulus parameters (Fig 3B). Studies by Dooling et al. (2001; budgerigars) and Lauer et al. (2006; budgerigars and humans) used the same tone frequency of 2.8 kHz as in the present study, whereas Kohlrausch and Sander (1995; humans) used tone frequencies of 2.2 and 4.4 kHz. In general, the budgerigar thresholds reported previously for F0 of 100 Hz (Dooling et al., 2001; Lauer et al., 2006) appear similar to those from the present study, showing the same V-shaped pattern with a minimum at C of approximately zero (Fig. 3B; blue symbols, left column). In contrast, prior budgerigar thresholds at 200 Hz (Fig. 3B, middle column) and 400 Hz (Fig. 3B, right column) are considerably higher and lower (Dooling et al., 2001), respectively, than those of the present study. Compared to previously published human results (Kohlrausch and Sander, 1995; Lauer et al., 2006) (Fig. 3B, red symbols), our findings suggest less threshold difference in budgerigars between Schroeder maskers with opposite C values.

Furthermore, budgerigars in the present study showed less increase in masked thresholds for higher F0 values than has been reported in humans, and intriguingly, appear substantially more sensitive than human subjects for F0 of 400 Hz and at all C values except +1 for F0 of 200 Hz.

#### 3.1.2. Roving-level paradigm

One difference between our study and previous studies of masking by Schroeder HTCs is that we used a roving-level paradigm for which a ±10 dB perturbation was applied to the stimulus level on each single-interval trial, while previous studies fixed the masker level, often at 80 dB SPL. A second experiment was conducted in four animals to evaluate the extent to which roving stimulus level impacted behavioral thresholds. C values of -1, zero, and +1 were tested using F0 of 100 Hz. The rove effect was assessed by transitioning from a fixed-level (80 dB SPL masker level) to roving-level test procedure (70-90 dB SPL masker level) partway through individual tracking sessions (Fig. 2B). Each animal completed the experiment over 2-3 months and 200-300 sessions.

Behavioral tracks showed decreasing tone level over the first 10-20 trials before stabilizing at the animal’s fixed-level thresholds, similar to roving-level tracks described above. Furthermore, most tracks showed no apparent change in tone level following the onset of the roving-level test period (Fig. 2B). Fixed-level masked thresholds calculated based on the final 25 unbiased tracking sessions (i.e., from portions of tracks prior to the rove onset) were lower for C of zero than for C of ±1 and similar between C values of -1 and 1 (Fig. 4A). These findings are consistent with results of the first behavioral experiment. Masked thresholds calculated from the roving-level part of tracks showed essentially the same pattern (Fig. 4B), suggesting minimal impact of this stimulus perturbation that makes single-channel energy cues less informative.

**Fig. 4.**
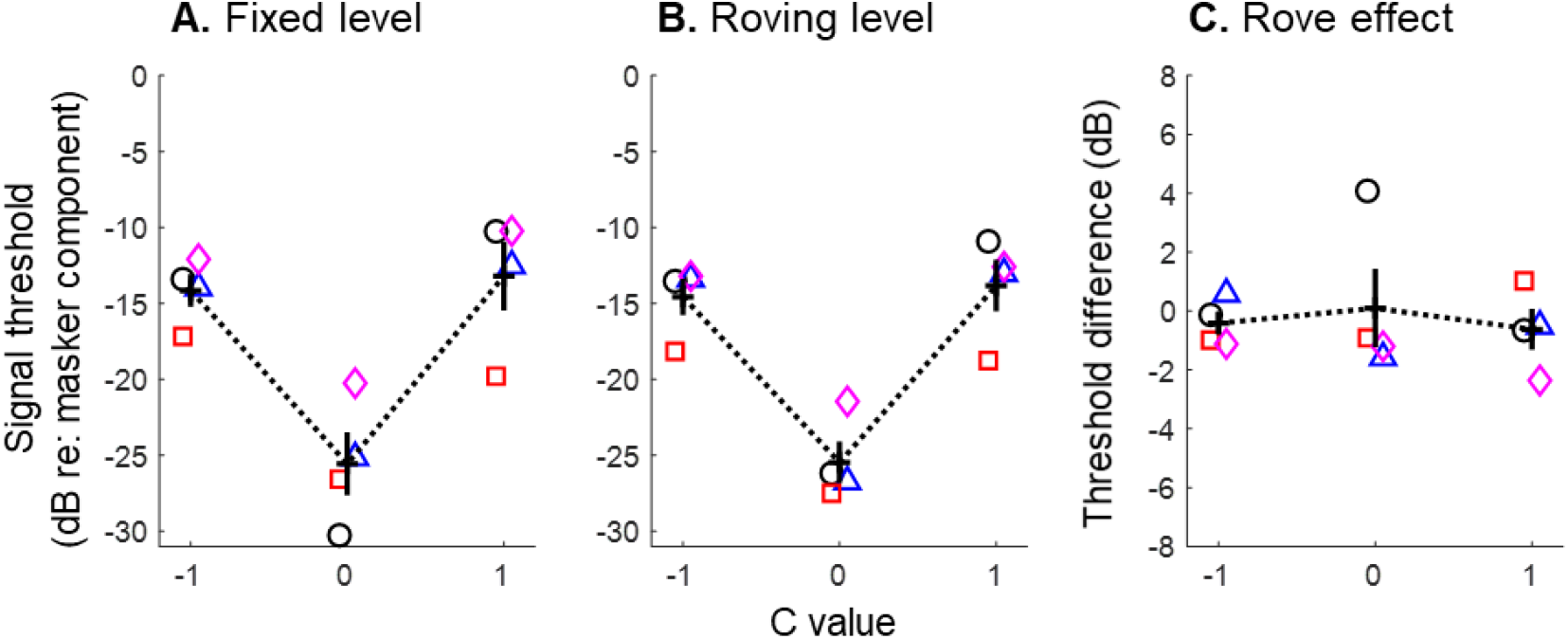
Behavioral Schroeder-masking thresholds of budgerigars from the second experiment comparing performance under fixed- and roving-level test conditions. Trend lines and error bars indicate means and cross-subject standard deviations, respectively. Symbols refer to animal identity as if Fig. 3. F0 is 100 Hz. Budgerigars showed little of no effect of roving stimulus level on Schroeder masking thresholds.

Finally, roving-level threshold shifts calculated from individual test sessions were generally near zero (Fig. 4C), though one animal (B34) showed a mean roving-level shift of 4 dB for C of zero only.

A mixed-effects model analysis of fixed- and roving-level thresholds showed a significant effect of C value (F_2,15_=69.57, p<0.0001) but no significant impact of roving level (F_1,15_=0.12, p=0.74) or the C value by roving-level interaction (F_2,15_=0.054, p=0.95). Thresholds were lower for C of zero than for C of -1 (−11.15 ±1.14 dB; t_15_=-9.82, p<0.0001) or +1 (−12.00 ±1.14 dB; t_15_=-10.57, p<0.0001; pairwise comparison of least-squared means ±1 SE), and similar between C values of -1 and +1 (−0.85 ±0.75 dB; t_15_=-0.75, p=0.47). The estimated effect of roving stimulus level was -0.32 ±0.93 dB (i.e., not significant; t_15_=-0.34, p=0.74). An ANOVA of within-track threshold shifts during the roving-level portion of tracks showed no significant intercept (−0.42 ±0.90 dB; t_8.9_=-0.46, p=0.65; i.e., mean roving-level shifts were not significantly different than zero) or effect of C (F_2,6_=0.18, p=0.84), consistent with results of the mixed-model analysis.

In summary, results from the two behavioral experiments support prior findings that Schroeder HTCs generated with positive and negative C values produce similar masking of a simultaneously presented tone signal in budgerigars. This result contrasts with prior human studies showing lower thresholds for Schroeder HTCs generated with positive C values (Kohlrausch and Sander, 1995; Lauer et al., 2006; Smith et al., 1986). Budgerigars also showed less apparent threshold elevation with increasing masker F0 than has been reported in humans. Finally, budgerigar thresholds were largely unaffected by the roving-level paradigm, suggesting that the animals used cues other than single-channel energy level to perform the masked-detection task.

### 3.2. Neurophysiological Results

#### 3.2.1. Neural frequency and amplitude-modulation tuning

Neurophysiological recordings were made from the budgerigar IC in awake animals (n=4; 142 multi-unit and 8 single-unit recording sites) to explore possible neural cues used by animals for Schroeder-masked tone detection. Detailed analyses of basic frequency tuning and modulation tuning in the budgerigar IC have been reported previously (Henry et al., 2017a; Henry et al., 2016; Wang et al., 2021). Briefly, frequency response maps of recorded IC neurons showed V-shaped excitatory tuning curves in response to tones (Fig. 5A), with thresholds at CF often falling below 20 dB SPL. CFs ranging from 0.54-5.6 kHz with a median value of 2.59 kHz.

**Fig. 5.**
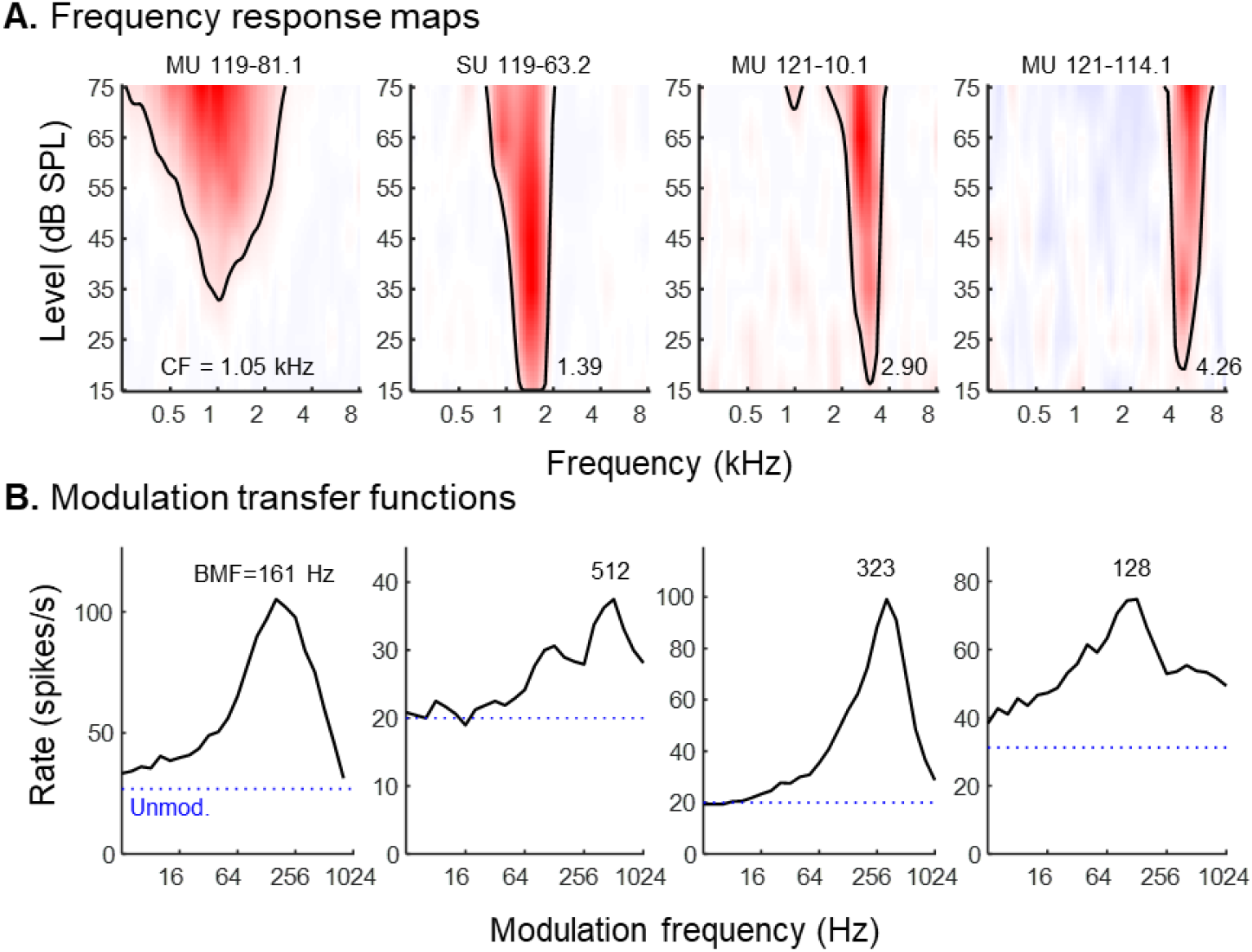
Basic tuning properties of recorded budgerigar IC neurons. Frequency response maps (A) show average rate as a function of tone frequency and level. SU: single unit; MU: multi unit; subsequent unit identification codes are arbitrary. Characteristic frequency (CF) increases from left to right. Modulation transfer functions (B) of the same recording sites in A, showing average response rate to sinusoidally amplitude-modulated tones across modulation frequencies. BMF: best modulation frequency. Dotted blue lines indicate the response rate to the unmodulated carrier signals. Carrier frequency is within ±1/6 octave of estimated CF.

An emergent response property at the IC processing level in birds and mammals is modulation tuning of average discharge rate in response to stimulus envelope fluctuations, typically quantified with SAM tones and a MTF approach (Fig. 5B) (Joris et al., 2004; Langner and Schreiner, 1988; Nelson and Carney, 2007; Woolley and Casseday, 2005). Most recorded budgerigar IC neurons exhibited some degree of band-enhanced modulation tuning (Kim et al., 2020) (i.e., 117 out of 125 units for which modulation transfer functions were obtained). That is, MTFs (Fig. 5B) showed greater response rate for a limited band of stimulus modulation frequencies compared to the unmodulated stimulus. Best modulation frequency (BMF), which is the modulation frequency associated with the peak rate of the MTF, ranged from 128-700 Hz across recorded neurons with a median value of 406 Hz. Neurons not showing band-enhanced modulation tuning had high-pass MTFs in all cases (n=8). The amplitude of modulation tuning, defined as the percent difference between the MTF peak and the response rate for the unmodulated stimulus (i.e., blue dotted lines in Fig. 5B), ranged from 22.5% to 199.8% with a median value of 120.5%.

#### 3.2.2. Effects of F0 and C on neural responses to Schroeder-phase HTCs without an added tone

Neurophysiological recordings made in response to Schroeder HTCs without an added tone showed robust fluctuations of discharge rate within F0 periods across the full range of F0 and C values studied (e.g., Figs 6A and 7A). Results from two representative IC units illustrate some of the diversity observed in Schroeder responses across neurons. Results from one neuron (Fig. 6) showed generally similar average rate and rate fluctuation (locked to F0 periodicity) between Schroeder HTC with opposite C values, but with moderate bias of rate fluctuation toward the positive Schroeder for F0 of 200 Hz and C values greater than approximately 0.5. In contrast, responses from a second neural recording (Fig. 7) showed pronounced bias toward the negative Schroeder, for both average rate and rate fluctuation, across most stimulus conditions tested.

**Fig. 6.**
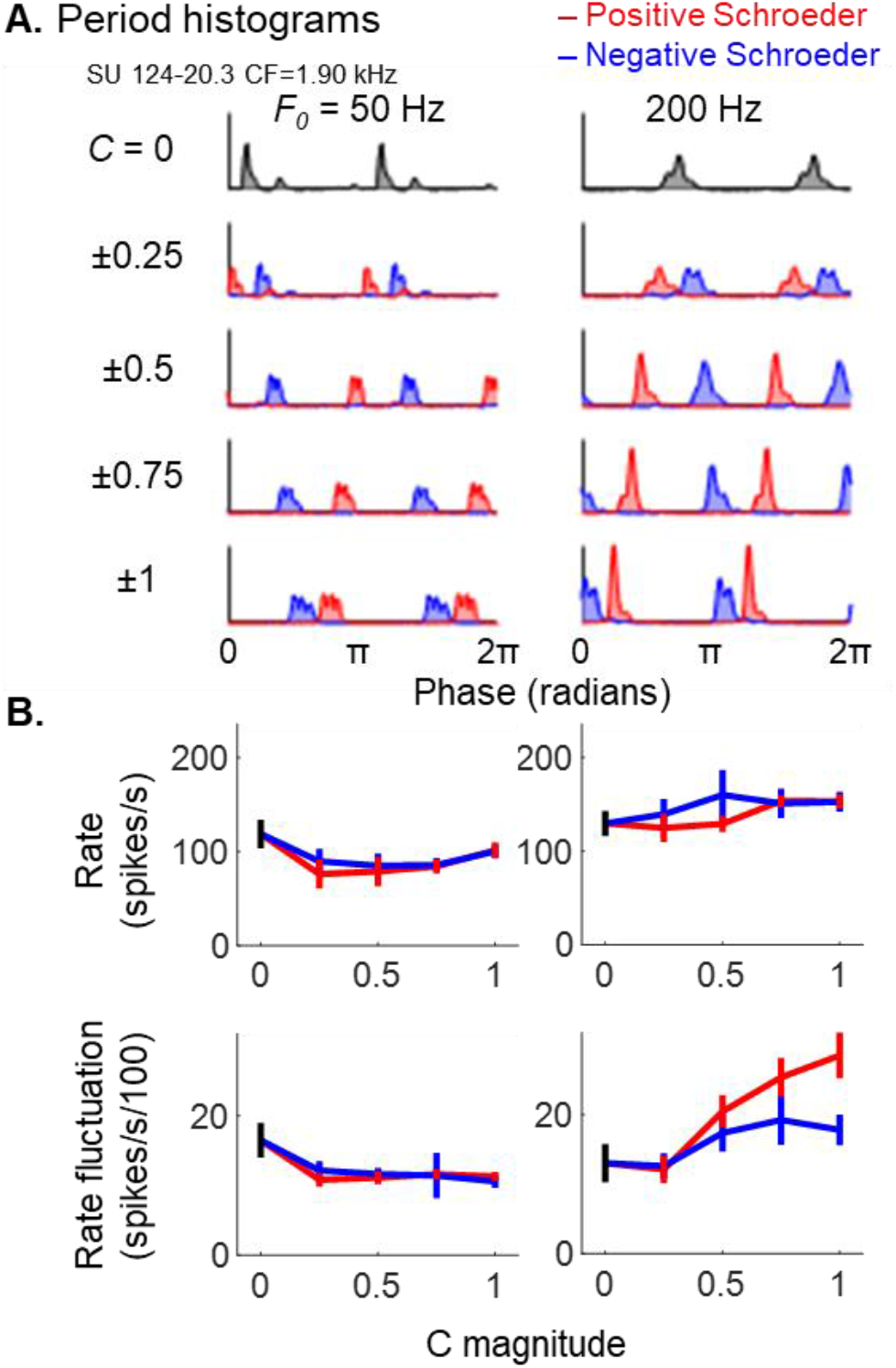
Representative neural responses to Schroeder-phase HTCs of variable F0 and C value. Period histograms (A) illustrate variation in average discharge rate over two cycles of F0. F0 is indicated at the top of each column and C magnitude given to the left. (B) Single-trial average discharge rate (top) and F0 rate fluctuation plotted as a function of C value for two different F0s (left: 50 Hz; right: 200 Hz). Trend lines and error bars show means and across-trial standard deviation. Note bias of F0 rate fluctuation toward the positive Schroeder for 200 Hz F0 and absolute C values greater than 0.5. CF was 1.90 kHz.

**Fig. 7.**
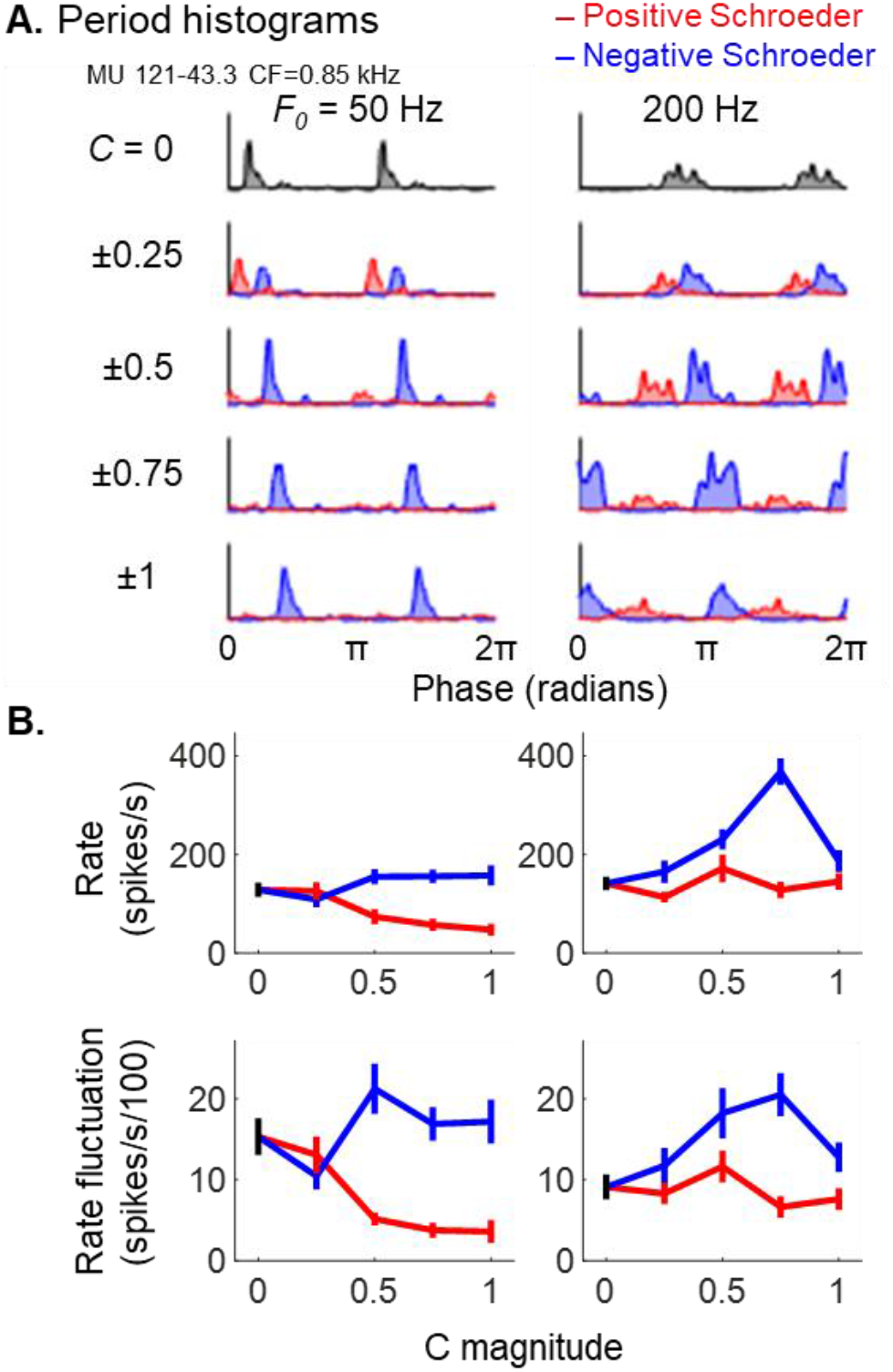
Representative neural responses as in Figure 5, but for a different unit. Note widespread bias of average discharge rate and F0 rate toward the negative Schroeder, most prominently for moderate-to-high absolute C values. CF was 0.85 kHz.

To characterize patterns of Schroeder response selectivity across the neural population, we calculated a Schroeder bias metric as the mean response difference between positive and negative Schroeder polarities divided by the pooled standard deviation. Positive (negative) bias values indicate greater response amplitude for the positive (negative) Schroeder polarity. Figure 8 shows Schroeder bias of rate fluctuation as a function of CF for each combination of stimulus F0 and C magnitude tested. Sample size was 146 for C of ±0.25 and ±0.75, and 150 for C of ±0.5 and ±1. Schroeder bias varied considerably across recorded IC neurons, ranging from negative to positive values for many conditions even among neurons of the same CF. Furthermore, mean Schroeder bias of the neural population increased with increasing CF, especially at high F0s and for higher absolute values of C (Fig. 8). The transition from negative to positive bias with increasing CF occurred at CF of ∼3 kHz. Schroeder bias of single units (Fig. 8, magenta circles) fell within the range observed for multi units of the same CF. Pearson correlations (Fig. 8; top left corner of each plot) showed that the relationship between CF and Schroeder bias was significantly positive for all stimulus conditions. Schroeder bias of average response rate followed the same general patterns reported above for rate fluctuation (Fig. 9).

**Fig. 8.**
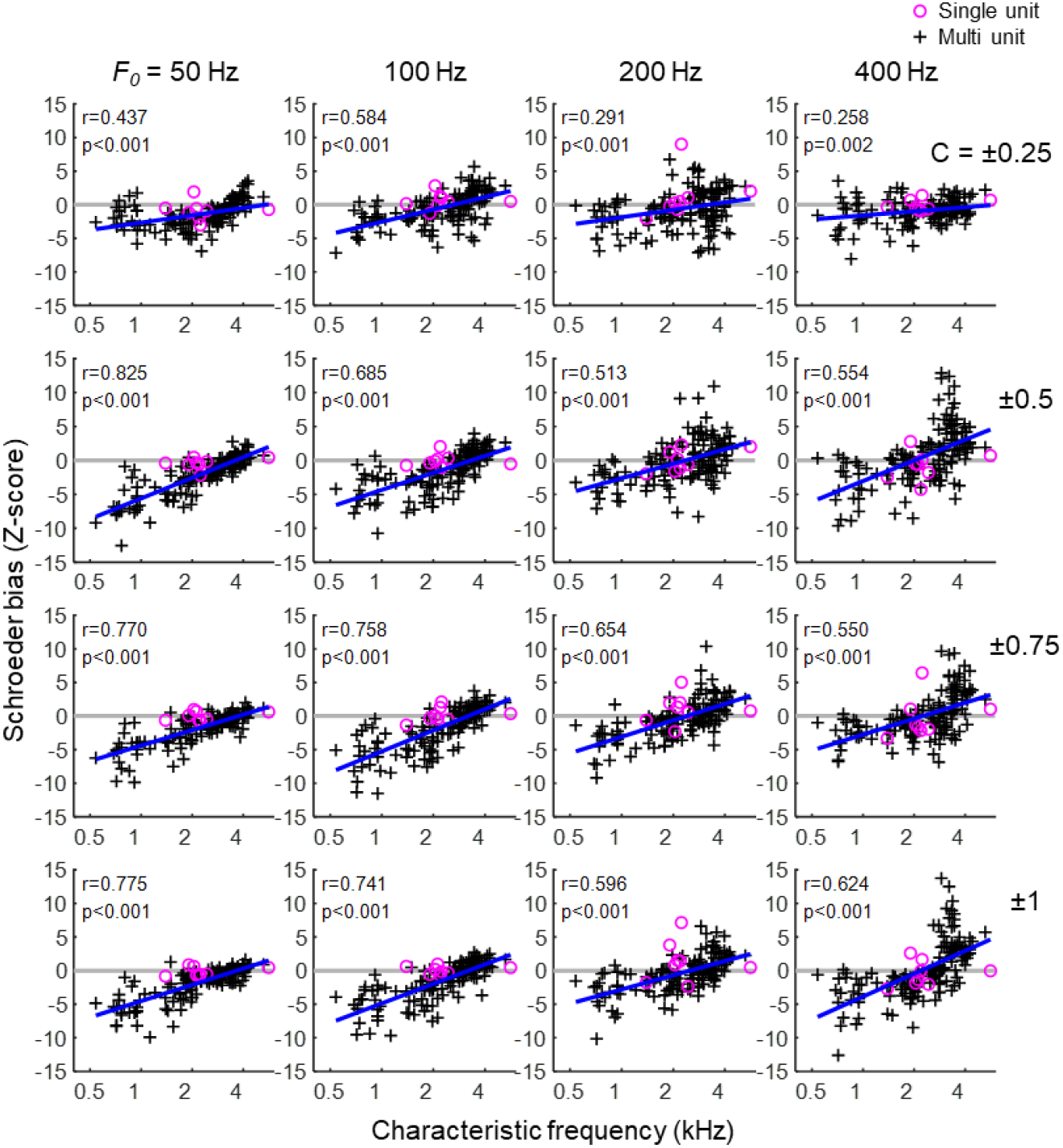
Schroeder bias of response rate fluctuation, plotted as a function of CF for all stimulus conditions. F0 is indicated at the top of each column and absolute C values are given on the right. Thick blue lines are predicted values from a linear fit to the data. Pearson correlations and associated p values are reported in the top left corner of each panel. The number of recorded units is 146 for C of ±0.25 and ±0.75, and 150 for C of ±0.5 and ±1.

**Fig. 9.**
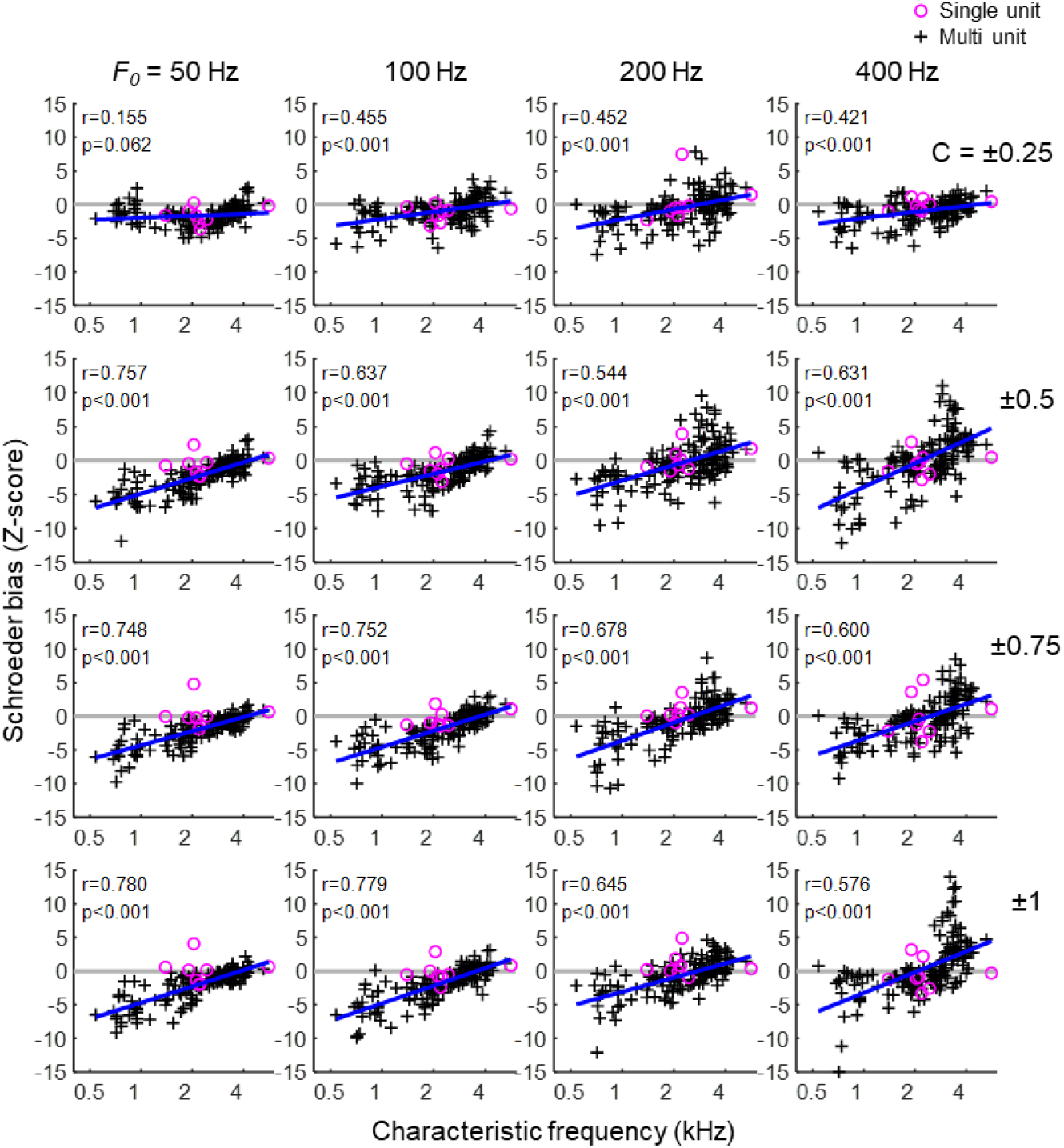
As for Fig. 8, but with Schroeder bias calculated based on average response rate.

#### 3.2.3. Neural thresholds for detection of Schroeder-masked tones

The changes in neural activity that could be used by behaving animals to detect tones masked by a Schroeder HTC were evaluated by recording IC neural responses to Schroeder-masked tone signals in a subset of units. Sample sizes were 92 for F0 of 100 Hz, 67 for F0 of 200 Hz, and 64 for F0 of 400 Hz. Masker level was 60 dB SPL and for each recording site, the signal frequency was set equal to that of the masker component nearest to CF. Thresholds were evaluated using ROC analyses, as the minimum signal level above which the area under the ROC curve (i.e., discrimination performance) consistently exceeded 70.7% correct. As illustrated in Fig. 10, many neural responses showed a pronounced reduction in activity as signal level increased. Neural threshold in these units was therefore based on a response decrement compared to the masker-alone condition (Fig. 10A, columns 1, 2, and 4; Fig. 10B).

**Fig. 10.**
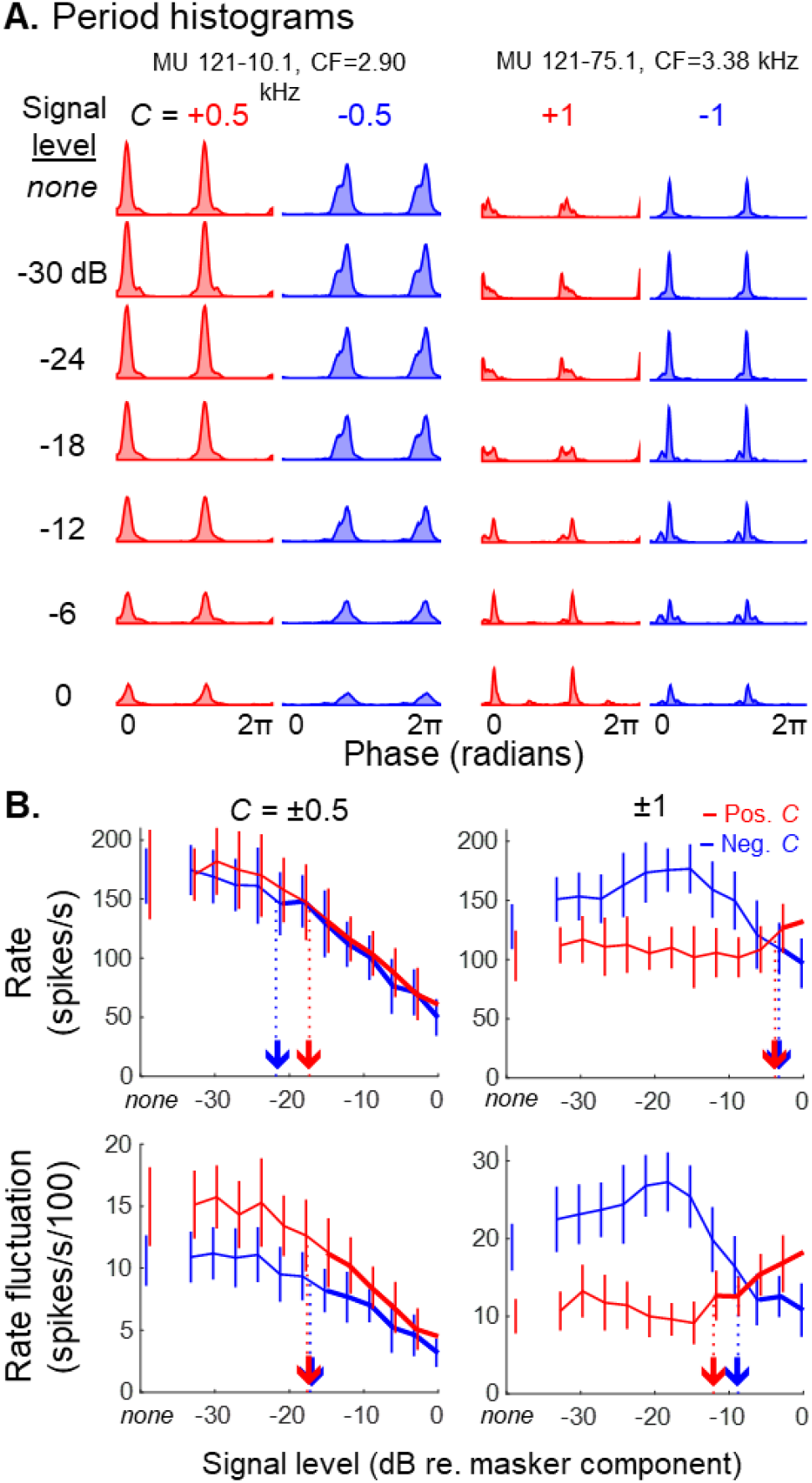
Representative neural responses to near-CF tones masked by Schroeder HTCs. Two-cycle period histograms (A) for a range of tone levels (indicated left) in two neurons; F0 is 400 Hz and C is ±0.5 (columns 1 and 2; first neuron) or ±1 (columns 3 and 4; second neuron). Response functions (B) show average discharge rate (top) and rate fluctuation (bottom) as a function of tone level. Downward pointing arrows indicate neural thresholds above which detection performance consistently exceeds 70.7% correct. Trend lines indicate cross-trial means and standard deviations. *None:* masker alone with no tone.

While perhaps counterintuitive, given that increasing signal level elevates the overall energy level of the stimulus, the finding is consistent with expectations for modulation-tuned IC neurons because increasing the signal level flattens the envelope of the peripheral response, and these neurons exhibit reduced response activity for sounds with weaker envelope fluctuations (Henry et al., 2017a; Henry et al., 2016; Krishna and Semple, 2000; Nelson and Carney, 2007). In contrast, detection thresholds of other neurons were based on response activity that increased with signal level (Fig. 10A, column 3; Fig. 10B, column 2, red), or non-monotonic response trajectories that elevated (Fig. 10A, column 4; Fig. 10B, column 2, blue) or precluded threshold estimation altogether. Note that different response profiles were sometimes observed in the same unit for opposite C values of the same masker F0 (e.g., Fig. 10A, columns 3 & 4; Fig. 10B, column 2).

Neural detection thresholds based on rate fluctuation are shown for the population of neurons as a function CF (Fig. 11). Among neurons with a measurable tone-detection threshold (29-82% of neurons dependent on condition), threshold was in most cases based on decreasing discharge rate consistent with IC modulation tuning (Fig. 11, blue circles and text) rather than a rate increase (Fig. 11; red triangles and text). Among neurons with CFs within ±¼ octave of the 2800-Hz signal frequency of behavioral experiments, thresholds increased for higher absolute C values (see medians and interquartile ranges in Fig. 11) and were similar between opposite polarities of the same C value, consistent with behavioral results (Fig. 11, large black circles). The mean value of measurable neural thresholds in this CF range was similar between negative and positive polarities of the same C value at F0s of 100 Hz (C ±1: 0.94 ±2.89 dB; t_23_=0.32, p=0.75; C ±0.5: 0.53 ±2.34 dB; t_25_=0.23, p=0.82), 200 Hz (C ±1: -2.84 ±2.67 dB; t_39_=-1.06, p=0.30; C ±0.5: 1.79 ±2.85 dB; t_50_=0.63, p=0.53), or 400 Hz (C ±1: -0.46 ±2.15 dB; t_37_=-0.21, p=0.83; C ±0.5: 2.10 ±2.17 dB; t_37_=0.97, p=0.34; mean threshold differences between negative and positive C value ±1 SE, and two-sample t-tests). On the other hand, whereas behavioral thresholds increased for higher F0s, neural thresholds showed the opposite pattern of lower values for higher F0s. Consequently, the most sensitive neural thresholds approximated behavioral thresholds for F0 of 100 Hz but were considerably more sensitive than behavioral performance for F0s of 100 and 200 Hz. Analyses of neural thresholds based on average rate produced the same conclusions (Fig. 12).

**Fig. 11.**
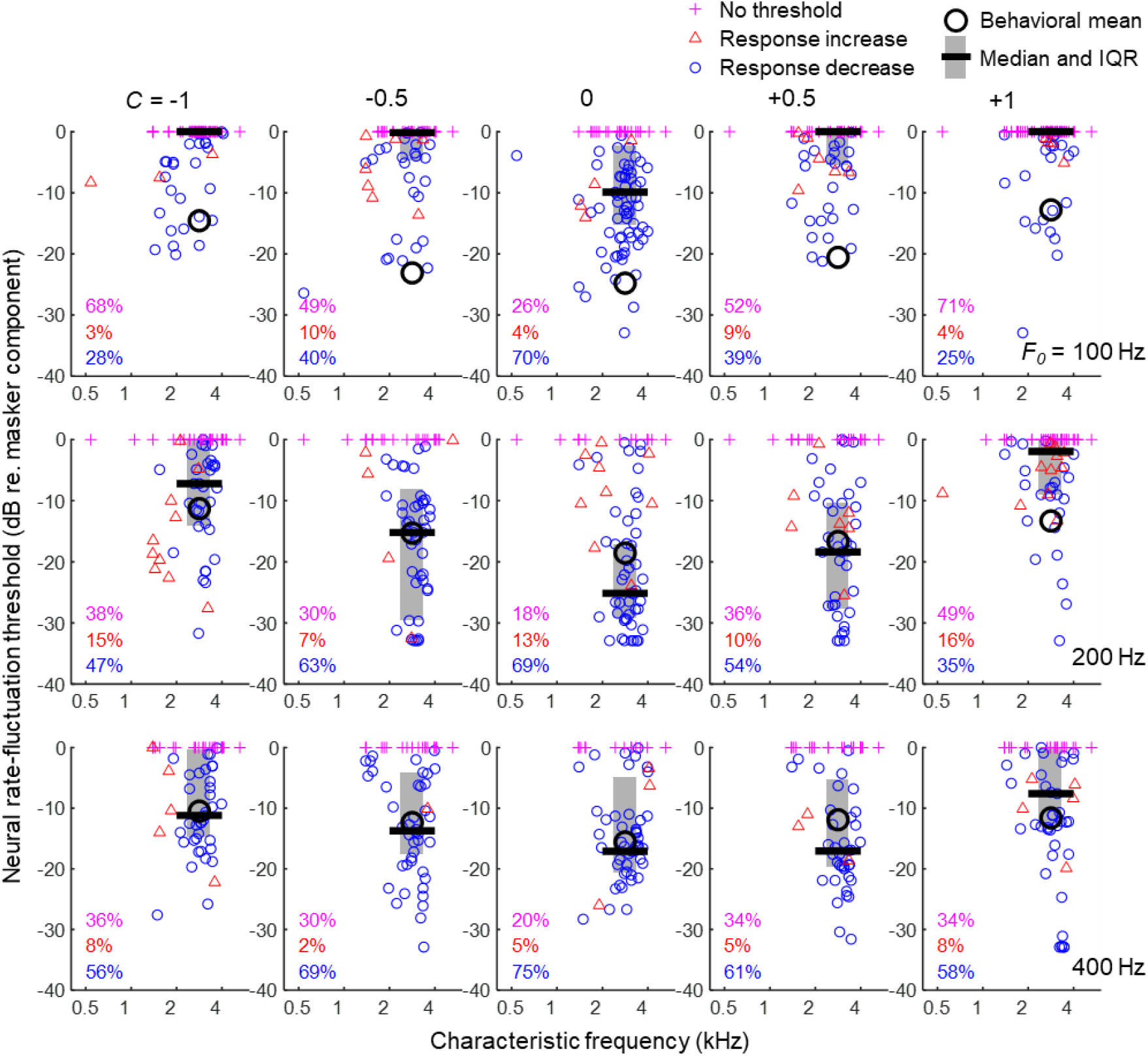
Neural thresholds for Schroder-masked tone detection based on rate fluctuation, plotted against CF. Tone frequency is the nearest harmonic frequency to CF and is within ±1/6 octave for all recordings. Masker level is 60 dB SPL. Threshold decreases for lower absolute C values, appears similar between opposite polarities of C, and increases for the lowest F0. Behavioral means are from Fig. 3. IQR: interquartile range. The total number of recorded units is 92 for F0 of 100 Hz, 67 for F0 of 200 Hz, and 64 for F0 of 400 Hz.

**Fig. 12.**
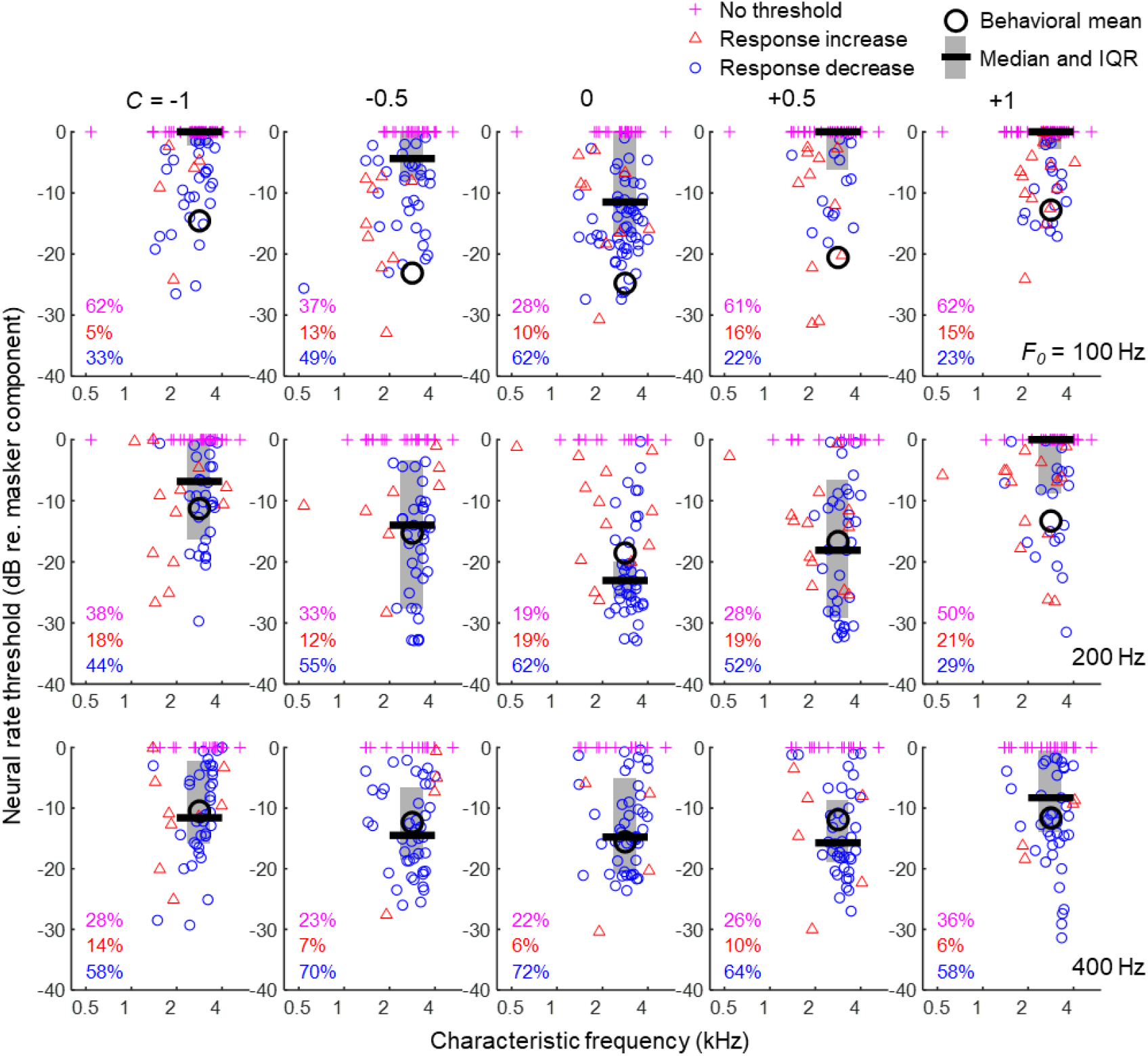
As for Fig. 11, but with neural thresholds calculated based on average rate. The total number of recorded units is 92 for F0 of 100 Hz, 67 for F0 of 200 Hz, and 64 for F0 of 400 Hz.

#### 3.2.4 Effects of Schroeder bias on neural masked-tone sensitivity

Finally, we evaluated the extent to which more extreme levels of Schroeder bias, as tended to occur in neurons with low and high CFs (see Fig. 8), might influence Schroeder-masked sensitivity to tones near CF. Separate analyses were conducted of signal thresholds assessed in Schroeder maskers with positive and negative C values. For F0s of 200 and 400 Hz, neurons with greater bias toward the positive Schroeder had lower tone-detection thresholds in the positive Schroeder masker (Fig. 13). Separation of neural data into low-, mid-, and high-CF ranges (Fig. 13, different symbols) further demonstrated that threshold variation was associated with the bias metric rather than CF per se. These results are broadly consistent with the premise from psychophysical studies that a peakier physiological response to the masker should enhance sensitivity to an added tone (Smith et al., 1986). On the other hand, neural thresholds for F0 of 100 Hz (both masker polarities; Fig. 13, top row), and for all maskers with negative C values (p=0.08-0.58), showed no significant association with Schroeder bias. Analyses of response metrics based on average rate produced the same conclusions (Fig. 14). Taken together, these results suggest that pronounced bias toward the positive-C Schroeder HTC can in fact enhance neural sensitivity for an added tone.

**Fig. 13.**
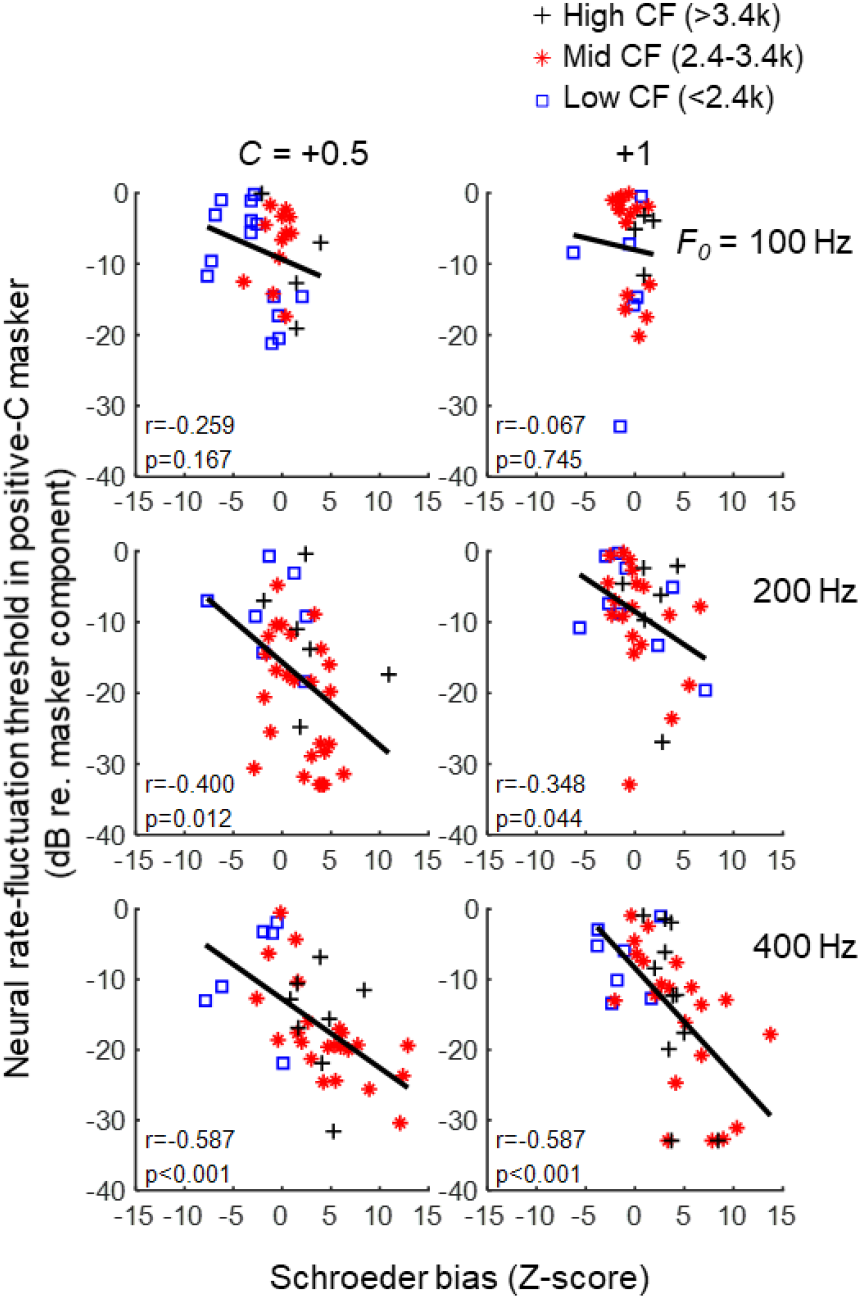
Neural thresholds for masked tone detection in positive Schroeder HTCs as a function of Schroeder bias observed for the same absolute C value. Both response metrics are based on rate fluctuation. Trend lines are a linear fit to the data excluding units with no threshold. Pearson correlations are reported in the lower left of each panel. Sample sizes are 30 and 26 for row 1 (left to right), 39 and 34 for row 2, and 37 and 39 for row 3. Greater positive-Schroeder bias is associated with lower masked thresholds for C values of +0.5 and +1 at F0s of 200 and 400 Hz.

**Fig. 14.**
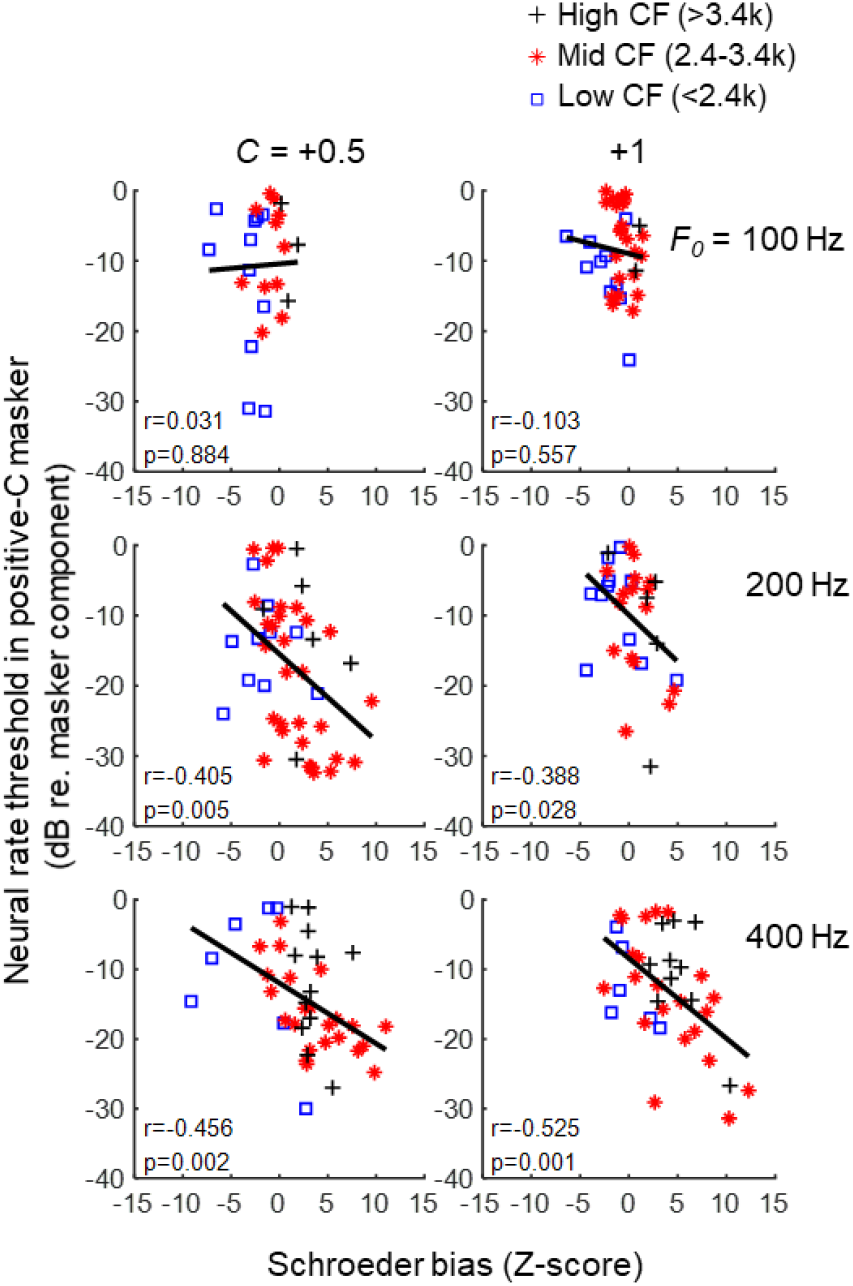
As for Fig. 13, but with neural thresholds and Schroeder bias calculated based on average rate. Sample sizes are 25 and 35 for row 1 (left to right), 46 and 32 for row 2, and 42 and 38 for row 3.

## 4. Discussion

Budgerigar behavioral thresholds for detection of a 2.8-kHz tone signal masked by a Schroeder-phase HTC were similar between C values of opposite polarity. Behavioral thresholds decreased for lower F0s, decreased for lower absolute C values, and were relatively unaffected by a roving-level paradigm that degrades the utility of single-channel energy cues for masked detection. Reponses of IC neurons to Schroeder HTCs showed considerable asymmetry between Schroeder HTCs with opposite C values in many cases, transitioning from average response bias toward the negative Schroeder at CFs less than 2-3 kHz to average response bias toward the positive Schroeder for higher CFs, but with substantial variability at any given CF. Neural thresholds for detection of Schroeder-masked tones near CF varied widely across IC units and were most often based on a response decrement upon addition of the tone signal, consistent with band-enhanced modulation tuning in IC neurons. Neural thresholds were generally similar between masker polarities, in agreement with behavioral results, though somewhat different effects of F0 were observed between behavioral and neurophysiological datasets. Finally, budgerigar IC neurons with pronounced bias toward the positive Schroeder HTC, as observed at high CFs, showed enhanced sensitivity to tones masked by positive Schroeder HTCs.

Behavioral results from the current study were largely consistent with prior studies of Schroeder masking in budgerigars, showing lowest thresholds for C values near zero (Lauer et al., 2006) and generally similar thresholds between Schroeder maskers of opposite polarity (Dooling et al., 2001; Lauer et al., 2006; Leek et al., 2000). That is, budgerigars showed symmetric V-shaped Schroeder masking as a function of C value. On the other hand, Dooling et al. (2001) found a different effect of masker F0 in budgerigars than was observed here, using C values of +1 and -1 and a fixed masker level of 80 dB SPL. Whereas thresholds in the current study increased monotonically with increasing F0, the earlier study found a substantial threshold reduction of 10-15 dB between F0s of 200 Hz and 400 Hz (Dooling et al., 2001). While the reason for this discrepancy is unknown, one difference between studies was that the current work used a roving-level paradigm to explore F0 effects, for which masker level varied randomly from 70-90 dB SPL across trials. The roving-level paradigm degrades the utility of single-channel energy cues for performing masked-detection tasks (Kidd et al., 1989; Richards, 1992). If animals in the prior fixed-level behavioral study relied more on single-channel energy cues for tone detection in maskers with F0 of 400 Hz,, disruption of the energy cue with the roving-level paradigm here could plausibly explain divergent F0 effects between our studies.

Schroeder-masked thresholds of human subjects can be 15-20 dB lower for maskers with positive C values compared to negative, depending on signal frequency, F0, and other factors (Kohlrausch and Sander, 1995; Lauer et al., 2006; Smith et al., 1986). These findings in humans are fundamentally different from the essentially symmetric V-shaped behavioral masking functions observed here, in prior studies of budgerigars, and in prior studies of other avian species including zebra finches and domestic canary (Dooling et al., 2001; Lauer et al., 2006; Leek et al., 2000). Unfortunately, behavioral studies of Schroeder masking have not yet been conducted in any nonhuman mammalian species, precluding comparison to our results. Compared to human results obtained with similar stimulus parameters to the present study, budgerigar thresholds at F0 of 100 Hz appear reasonably well matched to human data for C values from -1 to zero, but increase substantially between C of zero and +1 whereas human thresholds remain low across the same range (Kohlrausch and Sander, 1995; Lauer et al., 2006; Smith et al., 1986). Intriguingly, human studies have in fact found budgerigar-like V-shaped Schroeder-masking functions for signal frequencies of 250 Hz and below (Oxenham and Dau, 2001) and in listeners with sensorineural hearing loss (Summers and Leek, 1998). These results are perhaps attributable to reduced strength of fast-acting nonlinear cochlear gain (compression) at apical cochlear locations and in individuals with hearing loss.

Comparing masker-F0 effects on tone detection between species, while both budgerigars from the present study (but not Dooling et al., 2001) and humans show increasing thresholds for higher masker F0s, the differences are considerably more pronounced in the human data (Kohlrausch and Sander, 1995; Oxenham and Dau, 2001). For example, human thresholds for C of zero increase by ∼25 dB between F0s of 100 and 400 Hz (Kohlrausch and Sander, 1995), whereas budgerigar thresholds increased by just 10 dB over the same F0 range. Consequently, budgerigar behavioral thresholds were broadly similar to reported human data at the masker F0 of 100 Hz, excepting differences in masking asymmetry between C polarities, while being more sensitive than human results at 200 and especially 400 Hz. Budgerigars have also been found to discriminate between positive and negative Schroeder stimuli with greater accuracy than human subjects for stimulus F0s greater than or equal to 200 Hz (Dooling et al., 2002).

Neurons in the budgerigar IC often showed considerable response differences between Schroeder HTCs with opposite C values, in both average discharge rate and the amplitude of instantaneous-rate fluctuations within Schroeder F0 periods (i.e., referred to as “rate fluctuation” throughout). While variation was apparent within all CF ranges studied, both response metrics transitioned from bias toward negative C values at CFs below 2-3 kHz to bias toward positive C values at higher CFs. In other words, neural responses were stronger for HTCs with upward-sloping instantaneous-frequency sweeps at CFs below 2-3 kHz, but downward-sloping sweeps at higher CFs. These results could reflect a transition from downward-gliding to upward gliding impulse responses of the cochlea with increasing CF, consistent with what has been reported in several mammalian species (Carney et al., 1999; Recio-Spinoso et al., 2005; Recio and Rhode, 2000; Summers et al., 2003). On the other hand, it seems likely that “across-CF” combinations of excitatory and inhibitory inputs to IC neurons, with different timing, could produce neural sensitivity to sweep direction (Gittelman et al., 2009; Suga, 1965), or at least influence sweep sensitivity, potentially explaining differences in Schroeder bias observed between IC neurons of similar CF. Multiple excitatory inputs from auditory-nerve inputs with different CFs produces direction selectivity for frequency sweeps in octopus cells of the mammalian cochlear nucleus (Lu et al., 2022).

Differences in IC neural responses to Schroeder-phase HTCs with opposite C values have also been recently reported in anesthetized gerbils (Steenken et al., 2022). As in the present budgerigar study, Schroeder F0s were 50, 100, 200, or 400 Hz, and C values ranged from -1 to +1 with a step size of 0.25. Gerbil neural response differences were quantified using a discriminability index similar to our bias metric (but without polarity), based on average rate and temporal responses calculated using several window lengths, including 1/F0. Neural discriminability of Schroeder sweep direction was observed in most gerbil IC neurons based on average rate and, to a lesser extent, based on F0-related spike timing. Large response differences were frequently found between units, and these differences were not clearly related to CF or other basic response properties such as modulation tuning (see Fig. 4 in Steenken et al., 2022). Statistical analyses showed increasing Schroeder discriminability in gerbil IC neurons with band-reject modulation tuning, a response property not observed in our budgerigar study, and in neurons with higher BMFs (Steenken et al., 2022). Whereas average IC neural discriminability in gerbils was higher for low stimulus F0s, consistent with behavioral discriminability of Schroeder HTCs measured in the same species, individual IC neurons with high discriminability were found for every stimulus condition included in the study (Steenken et al., 2022). Taken together, these studies suggest that IC neural discriminability of Schroeder stimuli based on C polarity (i.e., direction of instantaneous-frequency sweeps) could be a broadly conserved aspect of complex-sound processing across diverse animal taxa.

Adding a tone signal to a Schroeder HTC increases the stimulus energy level, particularly in the processing channel centered on the signal frequency. Consequently, single-channel energy level is a potential cue by which human or animal subjects might detect the signal. Several lines of evidence in budgerigars argue against use of single-channel energy cues for Schroeder-masked tone detection. First, budgerigar Schroeder-masked thresholds, using a masker F0 of 100 Hz, were relatively unaffected by the roving-level paradigm that degrades single-channel energy cues. Similar results have been reported in prior behavioral studies of tone-in-noise detection in this species that compared fixed- and roving-level test paradigms (Henry and Abrams, 2021; Henry et al., 2020; Wang et al., 2021). Second, most IC neurons in budgerigars responded to Schroeder-masked tones in a manner consistent with modulation tuning and neural sensitivity to envelope fluctuations (Carney, 2018) rather than rate-based encoding of stimulus energy level. Neurons with band-enhanced modulation tuning, as commonly occur in the IC of many species including budgerigars, react to greater fluctuation depth of their inputs though increased average and synchronized response rate (Henry et al., 2016; Joris et al., 2004; Krishna and Semple, 2000; Langner and Schreiner, 1988; Nelson and Carney, 2007). IC inputs are highly modulated in response to the Schroeder masker alone, attributable to both the physical envelope of the HTC (for C values less than 1; Fig. 1) and the pronounced impact of band-pass filtering of the wideband HTC by the cochlea. These strongly fluctuating input signals become less modulated when the near-CF tone signal is added, resulting in a response decrement compared to the HTC masker alone due to modulation tuning.

While a response decrement could also feasibly result from inhibition by high-level tones, a previous study of budgerigar IC tone-in-noise sensitivity (Wang et al., 2021) suggests that this is not likely the case. Using tone signals added to third-octave narrowband noise, Wang and colleagues (2021) controlled level variation of both the signal and masker to help separate the roles of energy and envelope cues in determining IC response rates using a statistical model. Neural tone-in-noise thresholds were most often based on a response decrement compared to the masker alone, as in the present work on Schroeder masking. Multiple regression analyses demonstrated that response rate in most budgerigar IC neurons was positively correlated with the envelope cue, as expected based on IC modulation sensitivity, and was positively correlated with stimulus energy in contrast to expectations based on inhibition (Wang et al., 2021). Only a small percentage of units showed negative response correlations to stimulus energy, indicative of inhibition by high-level stimuli, most of which had CFs above 4 kHz, near the budgerigar’s upper frequency limit of hearing sensitivity. In summary, decrements in neural response amplitude appear to be a major cue for masked detection of pure-tone signals in the IC and likely result from the emergence of prominent modulation tuning at this processing level.

Budgerigar IC neural thresholds for Schroeder-masked tone detection showed several patterns broadly consistent with our behavioral results, including lower values for lower absolute C values and relatively similar values for opposite polarities of the same C value (i.e., V-shaped Schroeder masking functions). In contrast, whereas behavioral thresholds increased for higher masker F0s, neural thresholds showed the opposite pattern of lower values for higher masker F0s, regardless of whether they were calculated based on average rate or rate fluctuation.

While the reason for this difference is unknown, divergent results could feasibly reflect the fact that neural recordings were conducted with a relatively low, fixed masker level of 60 dB SPL, intended to minimize agitation of animals during awake, passive listening sessions. In contrast, the behavioral experiment exploring F0 effects was conducted with roving-level maskers of 70-90 dB SPL. If fixed- and roving-level thresholds increasingly diverge for higher masker F0s, this could explain discrepant F0 effects noted not only here, between behavioral and neural results, but also between behavioral results of the present study and those of Dooling et al. (2001) who use a fixed-level test strategy, as discussed previously.

While neural thresholds were on average similar between opposite C values, we found that units with strong response bias toward positive Schroeder HTCs were more sensitive to the addition of a tone presented with the positive Schroeder. This finding is consistent with general predictions in the Schroeder-masking literature (Smith et al., 1986; Summers and Leek, 1998). Because strong positive-Schroeder bias was most often observed at neural CFs above 3 kHz, for F0s of 200-400 Hz, and for absolute C value between 0.5 and 1, we would predict that behavioral Schroeder masking difference might exist in the budgerigar for these same conditions. Furthermore, we expect that thresholds should be lower for positive C values, as occurs in humans (Kohlrausch and Sander, 1995; Lauer et al., 2006; Oxenham and Dau, 2001; Smith et al., 1986). Unfortunately, behavioral testing in the present study was limited to 2.8 kHz, a signal frequency for which near-zero Schroeder bias was observed in IC neural responses. Furthermore, prior budgerigar studies that included a 4-kHz signal frequency were limited to a masker F0 of 100 Hz, for which little neural Schroeder bias was found in the present study. Future studies in budgerigars should test for asymmetry of Schroeder masking using 4-kHz tone signals and masker F0s from 200-400 Hz.

In summary, behavioral results of the present study showed V-shaped Schroeder-masking as function of C value in budgerigars, extending prior findings of this type in budgerigars and other bird species to a wider range of masking parameters. Schroeder-masked thresholds increased for higher masker F0s and were relatively unaffected by a roving-level paradigm, arguing against single-channel energy level as a major detection cue (at least for the 100-Hz masker F0). Neural recordings from the budgerigar IC in awake animals showed that tone detection in most neurons was based on a response decrement attributable to modulation tuning. This result highlights the likely importance of envelope cues and changes in the fluctuation strength of neural inputs (Carney, 2018) for masked detection. Neural thresholds showed symmetric V-shaped Schroeder masking functions, similar to behavioral results, but decreased somewhat for higher F0s in contrast to behavioral findings. Questions raised by the work include whether the roving-level effect, which was negligible behaviorally for a masker F0 of 100 Hz, might be more pronounced for higher masker F0s. Furthermore, future studies should determine whether asymmetric masking by Schroeder HTCs occurs behaviorally in this species for high signal frequencies (∼4 kHz) and when the masker F0 is 200-400 Hz.

